# UHRF1 suppresses viral mimicry through both DNA methylation-dependent and -independent mechanisms

**DOI:** 10.1101/2020.08.31.274894

**Authors:** RE Irwin, CA Scullion, SJ Thursby, ML Sun, A Thakur, SB Rothbart, GL Xu, CP Walsh

**Affiliations:** Genomic Medicine Research Group, Biomedical Sciences, Ulster University, Coleraine, BT52 1SA, UK; Institute of Biochemistry and Cell Biology, Chinese Academy of Sciences, Shanghai 200031, China; Laboratory of Medical Epigenetics, Institutes of Biomedical Sciences, Fudan University & Chinese Academy of Medical Sciences (RU069), Shanghai, China; Van Andel Research Institute, 333 Bostwick Ave, Grand Rapids, MI 49503, USA; Terry Fox Laboratory & Dept. of Medical Genetics, University of British Columbia, Vancouver V6T 1Z4, Canada

## Abstract

Some chemotherapeutic agents which cause loss of DNA methylation have been recently shown to induce a state of “viral mimicry” involving upregulation of endogenous retroviruses (ERV) and a subsequent innate immune response. This approach may be useful in combination with immune checkpoint cancer therapies, but relatively little is known about normal cellular control of ERV suppression. The UHRF1 protein can interact with the maintenance methylation protein DNMT1 and is known to play an important role in epigenetic control in the cell. To examine potential roles of this protein in differentiated cells, we first established stable knockdowns in normal human lung fibroblasts. While these knockdown cells showed the expected loss of DNA methylation genome-wide, transcriptional changes were instead dominated by a single response, namely activation of innate immune signalling, consistent with viral mimicry. We confirmed using mechanistic approaches that activation of interferons and interferon-stimulated genes involved in double-stranded RNA detection was crucial to the response. ERVs were demethylated and transcriptionally activated in UHRF1 knockdown cells. As in these normal cell lines, ERV activation and interferon response also occurred following the transient loss of UHRF1 in both melanoma and colon cancer cell lines. Restoring UHRF1 in either transient- or stable knockdown systems abrogated ERV reactivation and interferon response, but without substantial restoration of DNA methylation. Rescued cell lines were hypersensitive to depletion of SETDB1, implicating H3K9me3 as crucial to UHRF1-mediated repression in the absence of DNA methylation. Confirming this, cells rescued with UHRF1 containing point mutations affecting H3K9me3 binding could not mediate silencing of ERV transcription or the innate immune response. Finally, by introducing similar point mutations in the mouse homologue, we could show that this pathway is conserved in mice. Our results therefore implicate UHRF1 as a key regulator of ERV suppression and strengthen the basis for cancer cell hypomethylation therapy.

## Introduction

DNA methylation is known to play an important role in mice in maintaining suppression at many genes which are transcriptionally inactivated during development and differentiation (Smith and Meissner, 2013), such as those on the inactive X chromosome (Beard, Li and Jaenisch, 1995), silent alleles of imprinted genes (Li, Beard and Jaenisch, 1993), inactive olfactory receptor genes (McClintock, 2010), some protocadherins (Kawaguchi *et al*., 2008), and certain germline genes (Weber *et al*., 2007). These roles have largely been established by introducing mutations or deletions in the genes encoding the DNA methyltransferases either in the whole embryo, or in specific tissues such as the brain or germ line. DNA methylation has also been known for some time to be important for suppression of endogenous retroviruses in mice, as hypomorphic mutations in the maintenance methyltransferase DNMT1 result in widespread derepression of Intracisternal A Particles (*IAP*), a young and mobile class of ERV specific to rodents (Walsh, Chaillet and Bestor, 1998).

Less is known about the transcriptional response to the loss of DNA methylation in humans, where developmental models are lacking. Studies there have been hampered by a strong cell-autonomous DNA damage response which occurs even in undifferentiated cells lacking DNMT1, and acute loss of the enzyme results in cell death within a few cell generations through triggering a DNA damage response (Chen *et al*., 2007; Loughery *et al*., 2011; Liao *et al*., 2015). To circumvent this, we recently generated a hypomorphic series in human cells by selecting for the integration of an shRNA in a normosomic, untransformed normal lung fibroblast cell line hTERT-1604 (O’Neill *et al*., 2018). Here we found that chronic depletion of DNMT1 resulted in loss of methylation at some targets known from mice, such as protocadherins and olfactory genes, but also at some gene classes specific to human such as the cancer/testis antigen (CTA) genes. In fact, the transcriptional response in these cells was dominated by up-regulation of the CTA genes, the bulk of which are clustered on chromosomes X and Y (Simpson *et al*., 2005; Almeida *et al*., 2009).

Early work with pan-DNMT inhibitors (DNMTi) such as 5’-azacytidine (Aza) or 5’-aza-2’-deoxycytidine (5azadC) showed that CTA genes were one of the major direct targets of methylation-mediated repression in adult human tissues (Samlowski *et al*., 2005; James, Link and Karpf, 2006). These genes are normally expressed to varying levels in testis, but repressed elsewhere in the body in a methylation-dependent manner. Tumour cells often show genomewide hypomethylation and derepression of CTA genes, with presentation of fragments of these proteins on the cell surface (Karpf, 2006). More recent work with DNMTi in human tumour cell lines has shown that up-regulation of CTA genes is part of a cellular transcriptional response termed “viral mimicry” which appears to be triggered by loss of DNA methylation at endogenous retroviruses (ERV) (Chiappinelli *et al*., 2015; Roulois *et al*., 2015). In these studies, DNMTi treatment led to demethylation and transcriptional upregulation of ERV. The presence of double-stranded RNA (dsRNA) from ERV in the cytoplasm was recognised by the dsRNA sensors DDX58 (RIG1) and MDA5 (IFIH1), which triggered IRF7 signalling through the mitochondrial protein MAVS. IRF7 translocated to the nucleus and up-regulated interferons (IFN) and interferon-stimulated genes (ISG) which include dsRNA sensors and other upstream components in a feedback loop, triggering an innate immune response including presentation of antigens at the surface and cell-cell signalling (Chiappinelli *et al*., 2015; Roulois *et al*., 2015). The extent to which these effects are due to loss of DNA methylation only, or to secondary effects of the inhibitors is currently unclear, since viral mimicry has not been fully characterised in cells carrying DNMT1 mutations (Chiappinelli *et al*., 2015; Cai *et al*., 2017) and 5azadC is known to affect levels of the histone methyltransferase G9a (Wozniak *et al*., 2007), while Aza is mainly incorporated in RNA not DNA (Stresemann *et al*., 2006).

Mutations in the Ubiquitin-like with PHD and ring finger domains 1 (*Uhrf1*) gene (aka *Np95*) were initially characterised as phenocopying loss of DNMT1 in mouse and resulted in widespread hypomethylation of the genome and dysregulation of imprinted genes, as well as ERV such as *IAP* (Bostick *et al*., 2007; Sharif *et al*., 2007). In cells lacking UHRF1, the DNMT1 protein did not localise correctly to the nucleus and the paired tandem tudor (TTD)-plant homeodomain (PHD) region of UHRF1 has been proposed to allow interaction with chromatin even during mitosis by binding to histone 3 lysine 9 trimethylation (H3K9me3) (Rothbart *et al*., 2012, 2013). DNA methylation in mouse embryonic stem cells has been previously shown to have only some overlap with H3K9me3 deposited by the SETDB1 enzyme (Karimi *et al*., 2011). However, in post-implantation tissues, DNA methylation becomes more important even at regions marked by H3K9me3 (Wiznerowicz *et al*., 2007). Reports regarding the role of UHRF1, and more specifically H3K9me binding, in DNA methylation have varied. Mutations in the TTD-PHD region that affect H3K9me3 binding by UHRF1 have been shown in human to decrease DNA methylation at ribosomal DNA repeats in HeLa cells (Rothbart *et al*., 2012), but effects at single-copy genes and other regions of the human genome are unknown. In mouse, mutations in the same region gave only a 10% decrease in DNA methylation, which was genome-wide and not just restricted to repeats (Zhao *et al*., 2016). Other studies in mouse suggested that UHRF1 mutations caused widespread loss of methylation, but that it only played a minor role in ERV suppression (Sharif *et al*., 2016). In contrast, mutations in the zebrafish homologue were reported to result in ERV derepression in the developing embryo and activation of the innate immune system (Chernyavskaya *et al*., 2017) as for DNMTi in human, but through double-stranded DNA rather than dsRNA signalling.

There is, therefore, a lack of clarity regarding the role of UHRF1, what the cellular response to loss of this important epigenetic regulator would be, what genes would be most affected, and what the dependence, if any, of DNA methylation on the TTD-PHD domain would be. As complete ablation of UHRF1 caused cell death in mouse ES cells once differentiated (Bostick *et al*., 2007; Sharif *et al*., 2007), as well as in differentiated human cells (REI, MS, GLX, CPW data not shown), we used the same approach we recently took with DNMT1 (O’Neill *et al*., 2018) and generated a hypomorphic series using shRNA in a hTERT-immortalised normal fibroblast cell line as before. An unbiased genome-wide screen showed widespread loss of methylation across most regions, but the major transcriptional response was consistent with viral mimicry, including upregulation of innate immune and CTA genes. This appeared to be triggered by demethylation of ERV and the appearance of dsRNA in the cytoplasm. Rescuing the cells with intact UHRF1 could restore ERV repression and switch off the viral response. Interestingly, this occurred even in the absence of DNA methylation. Blocking H3K9me3-mediated silencing via knockdown of SETDB1 or mutation of the H3K9me3-binding pocket on UHRF1 prevented ERV suppression, suggesting this is upstream of DNA methylation. Consistent with this, the same binding pocket mutations cause loss of methylation at ERV in mice post-implantation, with concomitant derepression of ERV and innate immune genes. Our findings establish UHRF1 as a critical regulator of ERV suppression in mammals using a mechanism featuring H3K9me3 and DNA methylation.

## Results

### Generation of normal adult human cell lines depleted of UHRF1

We used our previously-described approach to generate cells lacking UHRF1 (Fig.1A). Briefly, normal human fibroblasts which have been immortalised using hTERT (hTERT-1604) were transfected with a construct containing shRNA and individual integrants selected for. Two rounds of experiments were carried out using shRNA targeting the main body (prefix U e.g. U5) or 3’UTR (prefix UH e.g. UH4, UH5) of the gene, and results were indistinguishable.

**Figure 1.**
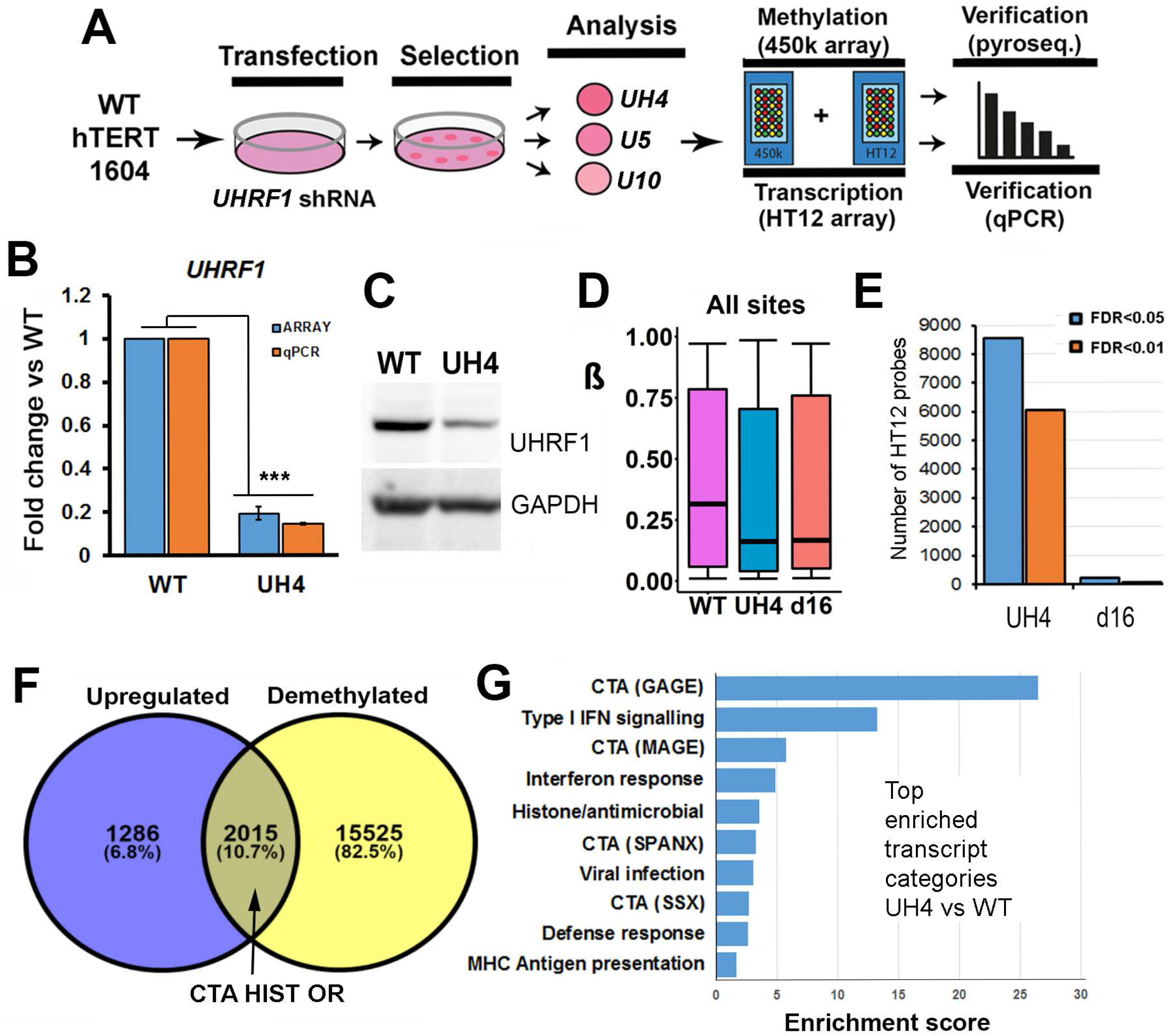
Establishment of human cell lines showing stable depletion of UHRF1 and genome-wide loss of methylation. (A)Schematic overview of the generation of UHRF1 knockdown cell lines: hTERT1604 normal fibroblasts were transfected with plasmid containing shRNA targeting the gene, as well as a selectable marker. Following growth in the presence of the antibiotic, surviving cells were expanded from single colonies and evaluated for genome-wide transcription and methylation using microarray. Results for individual loci were verified using pyrosequencing and RT-qPCR. (B) Comparison of HT12 transcriptional array and RT-qPCR results for UH4 (C) Western blot for UHRF1 (D) Overall median methylation from 450k array for *UHRF1*-depleted (UH4) cell line compared to the most demethylated *DNMT1*-depleted line (d16). (E) Greater numbers of probes from the HT12 transcriptional array showed significant differences between UH4 vs. WT than for d16 vs. WT. Numbers using different false discovery rates (FDR) are shown. (F) Genes which showed demethylation on the 450K array (n=18,826) were overlapped with those showing upregulation from the HT12 array. Gene names enriched in the area of overlap included those for cancer/testis antigens (CTA), histones (HIST) and olfactory receptors (OR). (G) The most up-regulated transcripts (n=3000) were analysed using DAVID functional annotation clustering: the top 10 enriched categories are shown. Different subclasses of CTA are shown in brackets; IFN, interferon; MHC, major histocompatibility complex. The x-axis represents group enrichment score, the geometric mean (in −log scale) of the p-values of the individual subcategories.

Those for the index line UH4 are shown here as an example. Results from other clones were similar and will be dealt with later (Suppl. Fig. 1A). Initial screening used reverse transcription-polymerase chain reaction (RT-qPCR, Suppl. Fig.1B). Cells showing depletion were further expanded, and UHRF1 mRNA and protein levels were checked by quantitative RT-PCR (RT-qPCR) and western blotting, respectively (Fig.1B,C). Lines showing depletion were further analysed using HT12 arrays for transcription, which verified low *UHRF1* levels (Fig.1B), as well as 450K arrays for DNA methylation (Fig.1D; Suppl. Fig. 1C). Median methylation levels, expressed as a □ value between 1 (fully methylated) and 0 (no methylation) were lower than WT in UH4 (Fig. 1D), and were comparable to the levels seen in our most severe hypomorph for *DNMT1* (d16), generated using shRNA in a similar manner and previously described in detail (O’Neill *et al*., 2018). Examination of the loss of methylation per genomic interval showed that UH4 had lower methylation in most regions, including CpG islands (CGI), gene bodies (genes) and in particular open sea, while d16 had greater loss overall at shores (data not shown). Notably, *UHRF1* hypomorphs with accompanying DNA hypomethylation were more readily isolated than DNMT1 hypomorphs, suggesting a more severe effect of the latter on cell viability.

### DNA demethylation in cells depleted of UHRF1 is accompanied by a specific innate immune response

Examination of the methylation profile across all promoter regions using 450K arrays indicated that loss of methylation was in general more severe in UH4 than d16 cells (Fig.1D). More probes on the HT12 transcription array also showed significant differences between UH4 and WT than for d16 versus WT (>8000 UH4 vs <500 d16 using a false discovery rate (FDR) of 0.05, Fig. 1E). DNA demethylation was widespread across the genome in UH4, with over half of all promoters (n=18,826), yellow circle (Fig.1F) showing significant (>0.1) decrease in β value. When these were compared with genes showing up-regulation from the set of dysregulated transcripts, the majority of genes (82.5%) showed demethylation but no derepression, which may reflect either no effect, or the absence of cell type-specific transcription factors required to activate them. A relatively small percentage (10.7%) were both demethylated and upregulated, consistent with a direct role for DNA methylation in their suppression in this cell type (Fig.1F). This included several previously characterised gene categories known to be regulated at least in part by DNA methylation, such as Cancer/Testis antigen (CTA) genes and olfactory receptors (OR), as we described previously in DNMT1 hypomorphs such as d16. Interestingly, a third group of genes (6.8%) showed no demethylation but were nevertheless up-regulated (Fig. 1F): this suggests an indirect response of genes in this category to loss of DNA methylation.

To investigate transcriptional response more closely, we then carried out gene ontology (GO) analysis of up-regulated transcripts from UH4 cells using the DAVID clustering tool (da, Sherman and Lempicki, 2009). Top hits in this analysis included several sub-classes of CTA genes (GAGE SPANX, MAGE). The other enriched gene categories included Type I interferon (IFN) signalling, antiviral response and MHC antigen presentation (Fig.1G). In fact, all 10 of the top 10 categories are related to the so-called “viral mimicry” state previously noted in cells treated with DNA methyltransferase inhibitors (Chiappinelli *et al*., 2015; Roulois *et al*., 2015).

### Innate immune signalling is crucial to the cellular response following loss of UHRF1

The viral mimicry state induced by methyltransferase inhibitors has been shown to be a response to the presence of dsRNA in the cell, which is detected by specific sensors in the cytoplasm (Fig.2A). These signal through the MAVS complex in the mitochondrion, releasing transcription factors (TFs) which turn on both interferons (IFN) and interferon-stimulated genes (ISG) in the nucleus (Jensen and Thomsen, 2012). Some of these latter gene products, such as the sensors DDX58 and OAS1 and the TF STAT1, are themselves components of the pathway leading to positive feedback-mediated amplification (broken arrows, Fig.2A). Consistent with this, a profile consisting of genes detected as up-regulated in our GO analysis, combined with previously reported genes, showed a clear up-regulation in UH4, but not d16 cells, compared to WT (Fig.2B). Profiling of the transcriptional response from the HT12 array (Fig.2C) showed clear up-regulation of components from several parts of the pathway shown in A. Changes in transcription level were most marked for ISG, including genes with anti-viral and cell death effects, and transcriptional changes were least marked or absent for TFs and MAVS (Fig.2C), as previously reported for this innate immune pathway (Cuellar *et al*., 2017). Notably, three of the genes unique to our profile and not previously reported are linked to T-cell signalling (Fig.2C). We verified sample genes from various parts of the pathway using RT-qPCR (Fig2D), with results consistent in direction, though not always in magnitude, between the array and the RT-qPCR. Of note, there was no evidence for up-regulation of components of the dsDNA response pathway from our array analysis, consistent with findings in cells exposed to DNA methyltransferase inhibitors.

**Figure 2.**
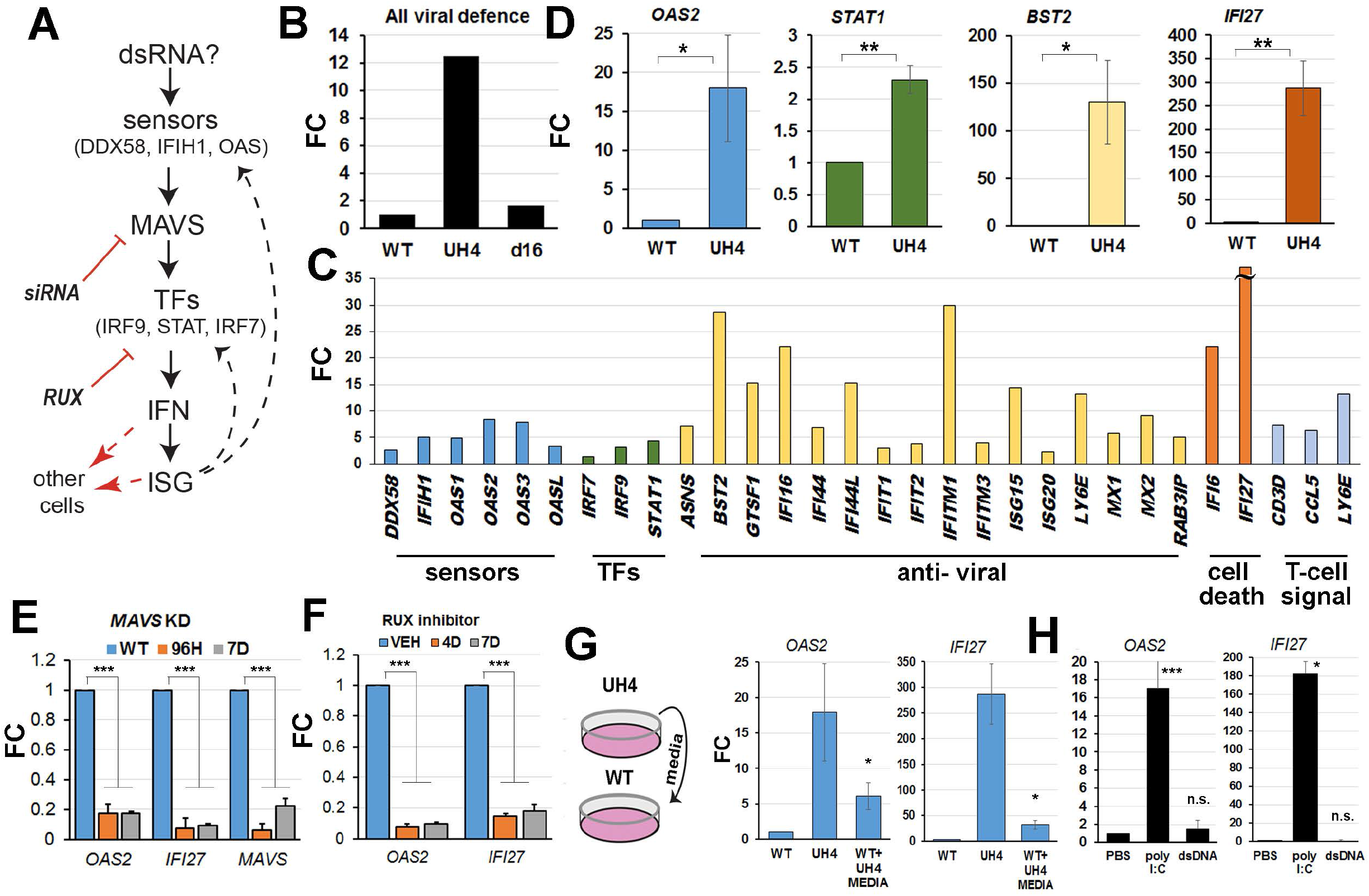
Interferons and interferon-stimulated genes (ISG) involved in dsRNA detection were crucial to the cellular response to loss of UHRF1. (A) Model for possible pathway triggering ISG response in UH4 based on GO analysis (Fig.2) and literature. Signalling from the dsRNA sensors would converge on the MAVS complex if dsRNA was detected, leading to activation of transcription factors (TFs) which trigger upregulation of both interferons (IFN) and ISG. Many ISG are also part of the pathway, leading to positive feedback (dashed arrows). Signalling to other cells can also occur (dashed red arrows). Inhibition of the pathway using siRNA against MAVS and the STAT inhibitor RUX are indicated. (B) Average fold change (FC) versus WT cells (set to 1) for viral defence genes from HT12 transcription array. (C) Many components of the signalling pathway are up-regulated at the transcriptional level. (D) Verification of selected targets from different parts of the pathway, using RT-qPCR. (E) An siRNA was used to knock down (KD) MAVS for the indicated period before assaying the indicated genes with RT-qPCR. (F) UH4 cells were treated with the JAK/STAT inhibitor RUX for the indicated time before carrying out RT-qPCR on the indicated genes. (G) Schematic of experiment where media which had been exposed to UH4 cells was transferred to WT cells (WT+UH4 media), before assaying gene transcription by RT-qPCR. (H) Exposure of WT cells to dsRNA (poly I:C) results in up-regulation of the same ISG as seen in UH4, measured here by RT-qPCR.

In order to investigate the dependence of cellular response on the activation of this innate immune pathway, we tested our model mechanistically. Inhibition of MAVS with siRNA in UH4 caused significant down-regulation of downstream ISG such as *IFI27* (Fig.2E). This included *OAS2* (Fig.2A), which although it is activated by dsRNA and therefore a sensor (Donovan *et al*., 2015), is also an ISG and up-regulated transcriptionally by anti-viral signalling (Fig.2C,D) in a feedback loop. Our GO analysis of transcriptional response in UH4 highlighted enrichment for genes involved in type I IFN signalling (Fig.1G), which included *IRF9* and *STAT1* (Fig.2C). Type I IFN binding at the cell surface activates JAK kinases, which phosphorylate STAT1 and STAT2, causing them to dimerise (Ivashkiv and Donlin, 2014). IRF9 associates with these dimers forming a complex termed the ISGF3 transcription factor which enters the nucleus and upregulates IFNs and ISGs (Fig.2A). We treated cells for 4-7d with Ruxolitinib (RUX), a small-molecule inhibitor of JAK kinases, and found a significant down-regulation of target ISGs, including *STAT1* itself (Fig.2F).

Our analysis so far suggested that components of the innate immune response are upregulated by depletion of *UHRF1* in the UH4 cells, including type I interferons (Fig.1G) as well as other cell surface and secreted signalling factors such as CCL5 and LY6E (Fig.2C). To test for cellcell signalling, we transferred media from tissue plates containing UH4 cells to plates with WT cells (Fig.2G): this resulted in up-regulation of ISG including *OAS2* and *IFI27*. All of the results above are consistent with an up-regulation of the dsRNA sensing pathway in the cells, presumably in response to the presence of dsRNA in the cytoplasm of UH4 cells (Fig.2A). Treatment of WT cells with polyI:C, a form of dsRNA, caused up-regulation of the same genes as seen in the UH4 line, confirming that the transcriptional response is consistent with exposure to dsRNA (Fig.2H).

### The presence of dsRNA correlates with transcriptional derepression and loss of DNA methylation at endogenous retroviruses in cells depleted of UHRF1

Type I interferon response can be triggered in cells when dsRNA is detected in the cytoplasm: this normally only occurs on infection of cells with viruses which produce dsRNA during their replication cycle, but can also occur if endogenous retroviruses are derepressed (Chiappinelli *et al*., 2015; Roulois *et al*., 2015; Cuellar *et al*., 2017). Staining of cells with the J2 monoclonal antibody is a sensitive and specific test for the presence of dsRNA (Weber *et al*., 2006) and gave a clear positive response in UH4, but not WT cells (Fig.3A). To test for derepression of ERVs, we used RT-qPCR for family members previously shown to be most active in response to epigenetic inhibitors (Cai *et al*., 2017; Cuellar *et al*., 2017), as ERVs are not covered on the HT12 array (Fig.3B). This indicated that members of several HERV families were transcriptionally up-regulated, including elements of the HERV-F (HERV-FC2), HERV-H (HERV-H) and HERV-W (HERV-W1) families (Fig.3C). As the fold change was small for a number of the HERVs, but J2 staining was much stronger than seen using polyI:C, suggesting the presence of large amounts of dsRNA, we considered that other ERV besides the HERV group might also be up-regulated in UH4. We therefore examined LINE-1 elements, another type of LTR-containing ERV element which can stimulate an IFN response and which are present at much higher copy number than HERV in the genome (Tie *et al*., 2018). RT-qPCR again found evidence consistent with up-regulation of some of these elements (*L1-PBA, L1P1*) in UH4 compared to WT controls (Fig.3D). Although small in magnitude (~2-fold), the absolute amount of dsRNA generated would be greatly increased due to the greater copy number of the elements involved.

**Figure 3.**
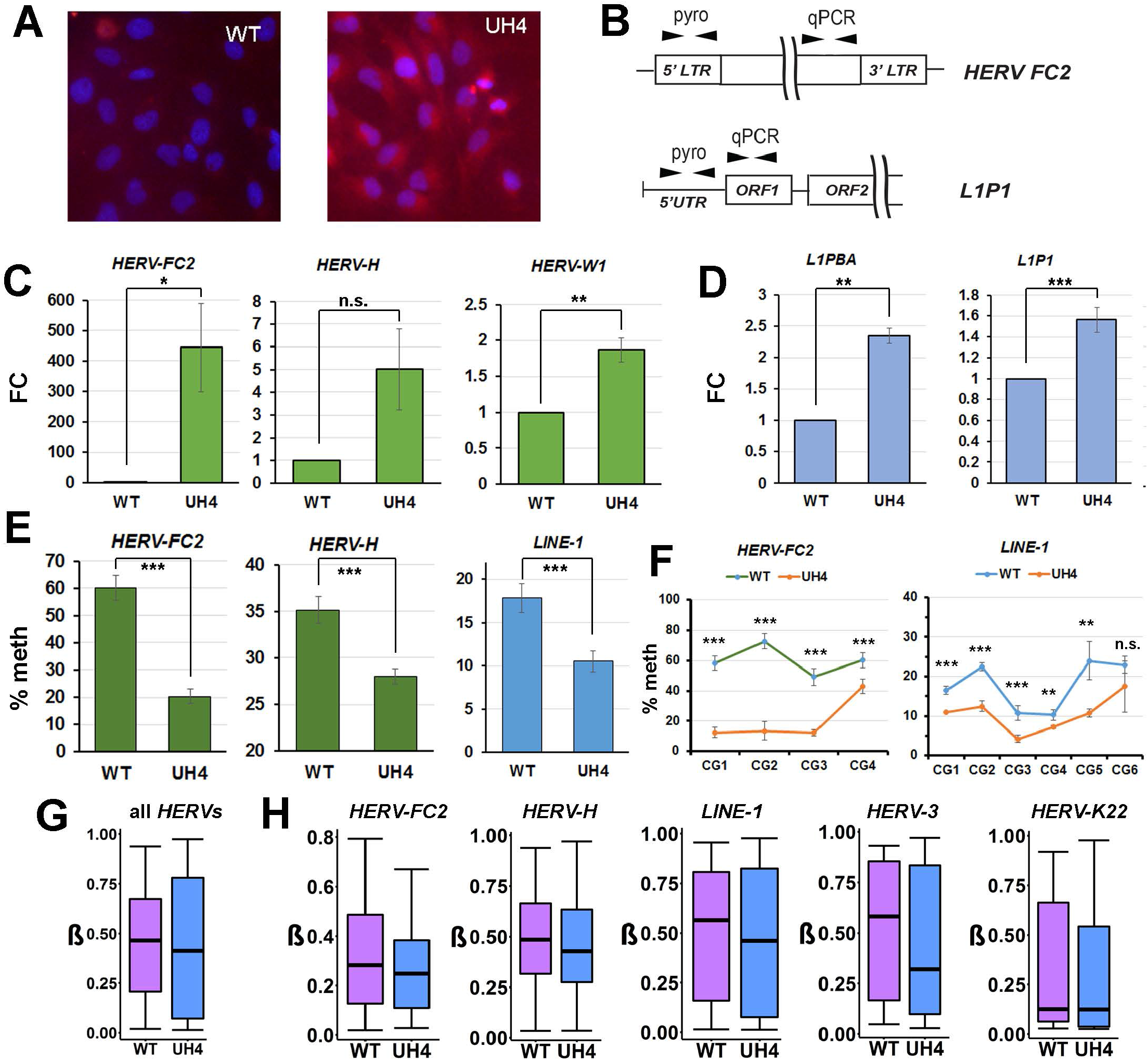
Human endogenous retroviruses (HERV) were demethylated and transcriptionally activated by depletion of UHRF1. (A) WT and UH4 cells were stained with J2 monoclonal antibody (red), used for detection of viral dsRNA; nuclei were counterstained with DAPI (blue). (B) Primer design locations for pyroassay and RT-qPCR (C) RT-qPCR for the indicated *HERV* elements showing fold-change over WT. (D) RT-qPCR for the *LINE-1* retroviral elements indicated. (E) DNA methylation (meth) at the 5’LTR of the indicated retroviral elements was determined using pyroassays (F) Methylation across individual CG dinucleotides in the pyroassays indicated in D. (G) Median methylation (β) at all probes overlapping HERV elements from the 450K array. (H) Median methylation at probes overlapping the indicated retroviral elements from the 450K array. Error bars indicate SEM (C,D,E,F) or 95% confidence interval (G,H).

Having established that specific elements were activated in UH4 cells, we examined control regions in these genes where methylation has been shown to act in a repressive capacity. Using pyrosequencing assays (pyroassays) covering multiple CG dinucleotides, we found consistent and significant demethylation of the ERV showing derepression, including *HERV-FC2, HERV-H* and *LINE-1* (Fig.3E). Examination of individual CGs in these regions showed significant demethylation across the entire region assayed (Fig.3F). While the 450K array was not designed to assay repetitive elements, a substantial number of probes overlap with regions labelled as ERV on the repeatmasker track in UCSC. Using the new bioinformatic workflow CandiMeth (Thursby *et al*. 2020), we assayed methylation across all ERVs in the genome, which showed a decrease in median methylation and greater variability in UH4 cells (Fig.3G). For regions with sufficient probe coverage, we also found evidence of substantial demethylation of several individual ERV families (*HERV-FC2, HERV-H, LINE-1, HERV-3*), but not for all (*HERV-K22*) (Fig.3G).

While the analyses so far have concentrated on the UH4 cell line, we confirmed our results in parallel for a number of other independently-derived clones from the two rounds of transfection (Suppl. Fig.2B,C).

### A conserved interferon response follows ERV demethylation in multiple cell types

Our results so far strongly supported a role for UHRF1 in methylation and repression of ERV in the hTERT-1604 normal fibroblast line and showed that stable depletion resulted in a robust innate immune response targeted against the viral RNA. We then wished to examine the timing of these events, both in the non-transformed hTERT1604 and in different transformed cells to determine whether loss of UHRF1 triggers the same transcriptional response. Further, we wished to determine whether cells could recover from the loss of the protein and re-establish repression. To this end, we carried out a transient or “hit-and-run” experiment (Fig.4A) where we exposed cells to siRNA against *UHRF1* for 48hrs, then switched the cells to normal medium without siRNA and allowed them to recover for up to four weeks. RT-qPCR showed that *UHRF1* levels were effectively depleted to ~25% by 96hrs, after which point they steadily recovered, reaching 50% by 14 days (14D), and continuing to rise up to and beyond (1.5 fold higher) levels seen in scrambled controls (SCR) by 28D (Fig. 4B). Consistent with observations in our stable knockdown clones, *HERV-H* mRNA levels were increased versus scrambled controls, starting already at 96hrs, and climbed steadily until 14D, at which point they started to decrease and were back at levels seen in SCR control by 28D (Fig.4B). The ISG gene IFI27 showed a similar dynamic, increasing from 7D and then decreasing to normal levels or below by 28D. Examination of methylation levels at ERV by pyroassay showed loss of methylation at the promoter regions of ERVs already at 72hr (Fig.4C). Interestingly, methylation showed only a moderate gain during the recovery period and remained significantly lower than WT out to 36D, beyond the period during which transcription of the ERV and ISG had already normalised, for both average methylation and levels at individual sites across the promoters (Fig.4C). Non-recovery of methylation was evident at both 7D and 21D for *LINE-1* (Fig.4D).

**Figure 4.**
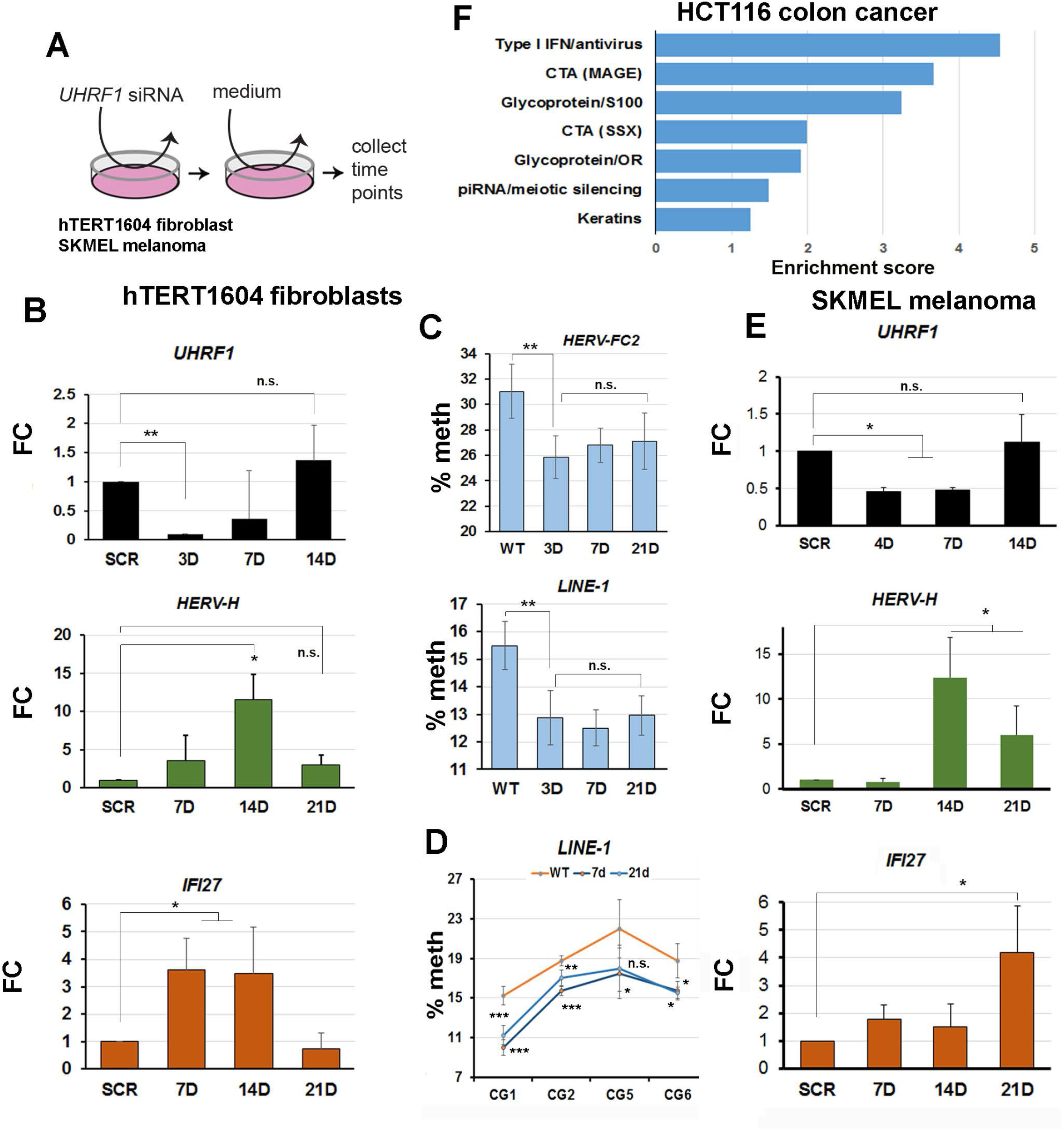
Demethylation of HERVs precedes reactivation and an interferon response in multiple cell types depleted of *UHRF1*. (A) A “hit-and-run” strategy was employed to establish timing of events: indicated cell types were exposed to UHRF1 siRNA for 48hrs, then fresh medium without siRNA added and cells allow to recover for the times indicated below before sampling. (B) RT-qPCR showing initial loss of *UHRF1* is followed by recovery to above initial levels by 28 days. Levels of transcript for a representative ERV (*HERV-H*) and ISG (*IFI27*) are shown. FC, fold-change. (C,D) Average methylation levels at representative retroviral elements as determined by pyroassay (top); error bars are SD. Methylation recovery by 21days, when transcription is already repressed, is still insignificant (bottom). The most highly-methylated sites are shown, effects are similar at CG3 and CG4; error bars are SD. (E) RT-qPCR analysis of SKMEL melanoma cells treated as in B. (F) GO analysis of genes showing transcriptional upregulation in HCT116 colon cancer cells following 90% KD of *UHRF1* (Cai et al, 2018). Enrichment scores etc as above. Cells were analysed 7days after adenoviral delivery of shRNA; raw data were obtained from GEO (GSE93136).

We then sought to determine if similar transcriptional responses would be seen in tumour cells. To this end, we performed a similar transient KD and recovery experiment in SK-MEL-28 melanoma cell lines, which have a more epithelial character. While transient KD was less efficient in these cells, UHRF1 levels were depleted to ~50% by 7D, then rapidly recovered to levels seen in scrambled controls by 10D (Fig.4E). This was accompanied by delayed activation of ERV and ISG by 10D, peaking at 14D after which point transcription of both started to decrease again (Fig.4E).

Additionally, we reanalysed a publicly-available dataset (Cai *et al*., 2017) where *UHRF1* was depleted in HCT116 colon cancer cells using adenovirus-mediated transfection of shRNA and where some up-regulation of ISG had been reported. Using GO analysis, we found that the top enriched gene class was Type I interferon response, with CTA activation accounting for two more of the top 5 categories (Fig.4F). An additional category among the top 7 was piRNA/meiotic silencing (Fig.4F). Taken together with the results above, this suggested that UHRF1 depletion leads to reproducible ERV demethylation and derepression in multiple cell types, evoking a strong innate immune response.

### Rescuing stable KD clones with UHRF1 can restore ERV repression without reestablishing normal DNA methylation levels

The transient experiments also suggested, importantly, that ERV repression could be reestablished without full remethylation. In order to confirm this in a more stable system, we undertook to rescue UHRF1 expression in UH4 cells by transfecting them with full-length cDNA lacking the 3’UTR which is targeted by the shRNA in UH4 but containing a FLAG epitope (Fig.5A). Western blotting confirmed the presence of the full-length, FLAG-tagged protein in rescues, termed WT10 (Fig.5A). Despite DNA methylation remaining at UH4 levels, the WT10 cells showed clear restoration of repression at HERVs (HERV-FC2, HERV-H) and LINE-1 elements (L1PBA) (Fig.5B). Normalisation of ISG levels was also seen in WT10 cells by RT-qPCR (Fig.5C). Analysis of overall transcription by HT12 array confirmed widespread shut-down of the innate immune response, with genes from most components of the pathway returning to normal or near-normal levels (Fig.5D, black columns), with the exception of a few genes (GTSF1, BST2). Examination of the methylation levels using 450k arrays showed that, despite the presence of WT UHRF1, median methylation levels in WT10 were indistinguishable from UH4 (Fig.5E). There was no increase in median methylation from UH4 over HERVs as a whole, or in the HERV-FC2 subclass, as confirmed by both array and pyroassay analysis (Fig.5F,G). The same was true of LINE-1 elements, where methylation at the promoter showed no change either (Fig.5F,G). These results, taken together with the transient experiment (Fig.5), indicate that UHRF1 can restore repression even when DNA methylation levels cannot be fully re-established.

**Figure 5.**
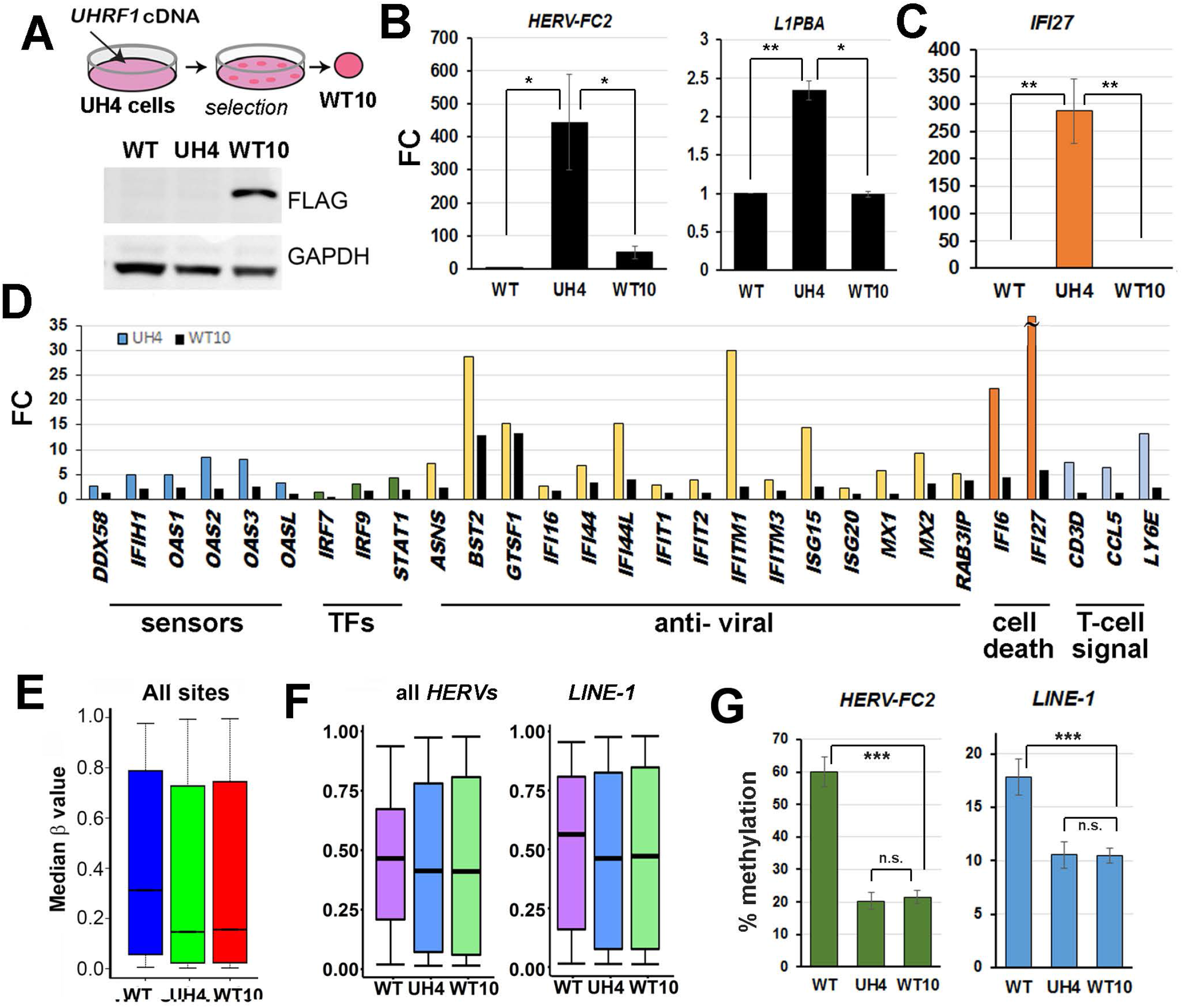
Rescuing cells with UHRF1 could abrogate HERV reactivation and interferon response without restoring DNA methylation. (A) Schematic (top) showing rescue strategy: a plasmid containing a selectable marker and a full-length *UHRF1* cDNA lacking the 3’UTR but with a FLAG-tag was transfected into UH4 cells and resistant colonies expanded. Western blot (bottom) detected the presence of the fulllength FLAG-tagged UHRF1 in daughter cell line WT10. (B) RT-qPCR of individual ERVs shows re-establishment of repression in WT10 cells; error bars represent SEM. (C) Selected ISG also show shut-down in WT10 cells expressing full-length UHRF1 by RT-qPCR. (D) HT12 array results for WT10 show most ISG involved in the response pathway are down-regulated again with 1-2 exceptions (e.g. *BST2, GTSF1*). (E) 450K analysis indicated that median methylation (β) levels were largely unchanged in WT10 cells; error bars represent confidence intervals (F) Methylation across all *HERV* and *LINE-1* elements assessed by 450K (G) Median methylation at *HERV-FC2* and *LINE-1* assessed by pyroassay, error bars are SEM.

### Hypomethylated cell lines rescued using mutated UHRF1 proteins implicate the KAP1/SETDB1 system in ERV repression

To explore what other mechanisms may be engaged by UHRF1 to repress ERV in the absence of DNA methylation in WT10 and transient KD cells, we considered the KAP1/SETDB1 system. The strong transcriptional corepressor KAP1 has been shown in mouse (Wiznerowicz *et al*., 2007), and only very recently in human (Tie *et al*., 2018), to repress ERV and to cause the deposition of both DNA methylation and H3K9 trimethylation (H3K9me3) on the silenced elements, the latter through the histone methyltransferase SETDB1 (Fig.6A). Since WT10 rescue cells represent a stable system with low DNA methylation on ERV, we reasoned that these may show greater sensitivity to loss of SETDB1 if these proteins were important for retroviral repression. Transfection of both normal hTERT1604 fibroblast cells (WT) and the rescue cell line lacking DNA methylation (WT10) with siRNA targeting SETDB1 gave a marked derepression of ERV and activation of ISG in WT cells, and an even greater response in the WT10 cells (data not shown). SETDB1 KD showed marked decrease in H3K9me3 marks at the protein level in WT10 cells (Fig. 6B). The UHRF1 protein has been previously shown, through both crystallographic and binding studies, to engage the H3K9me3 mark through its paired tandem Tudor domain and plant homeodomain (TTD-PHD), with key residues including Y188 (TTD) and D334/E335 (PHD) (Rothbart *et al*., 2012, 2013) (Fig. 6C). We used the same constructs as before to rescue UH4 cells and isolate clones expressing FLAG-tagged UHRF1 proteins containing these mutations in either the TTD (TTD9) or PHD (PHD1, PHD4, PHD10) domains (Fig.6D), which expressed the rescued protein to various extents (Fig.6E). As for WT10, none of the rescues re-established methylation at ERV elements (data not shown), but unlike cells rescued with intact protein (WT10, WT18), the cell lines containing mutated UHRF1 showed poor and variable repression of ERV (Fig.6F). Furthermore, cells with the point mutations were positive for dsRNA in the cytoplasm using J2 staining (Fig.6G). In this respect, they resembled cells with little UHRF1 (UH4), whereas cells rescued with WT protein (WT18) showed little or no staining (Fig.6G). In keeping with the failure to repress ERV, there remained a robust ISG response (Fig.6H) in the UH4 cell lines rescued with the point-mutated UHRF1 (PHD1, PHD4, PHD10, TTD9), but not when the same UH4 cells were rescued with intact protein (WT10, WT18).

**Figure 6.**
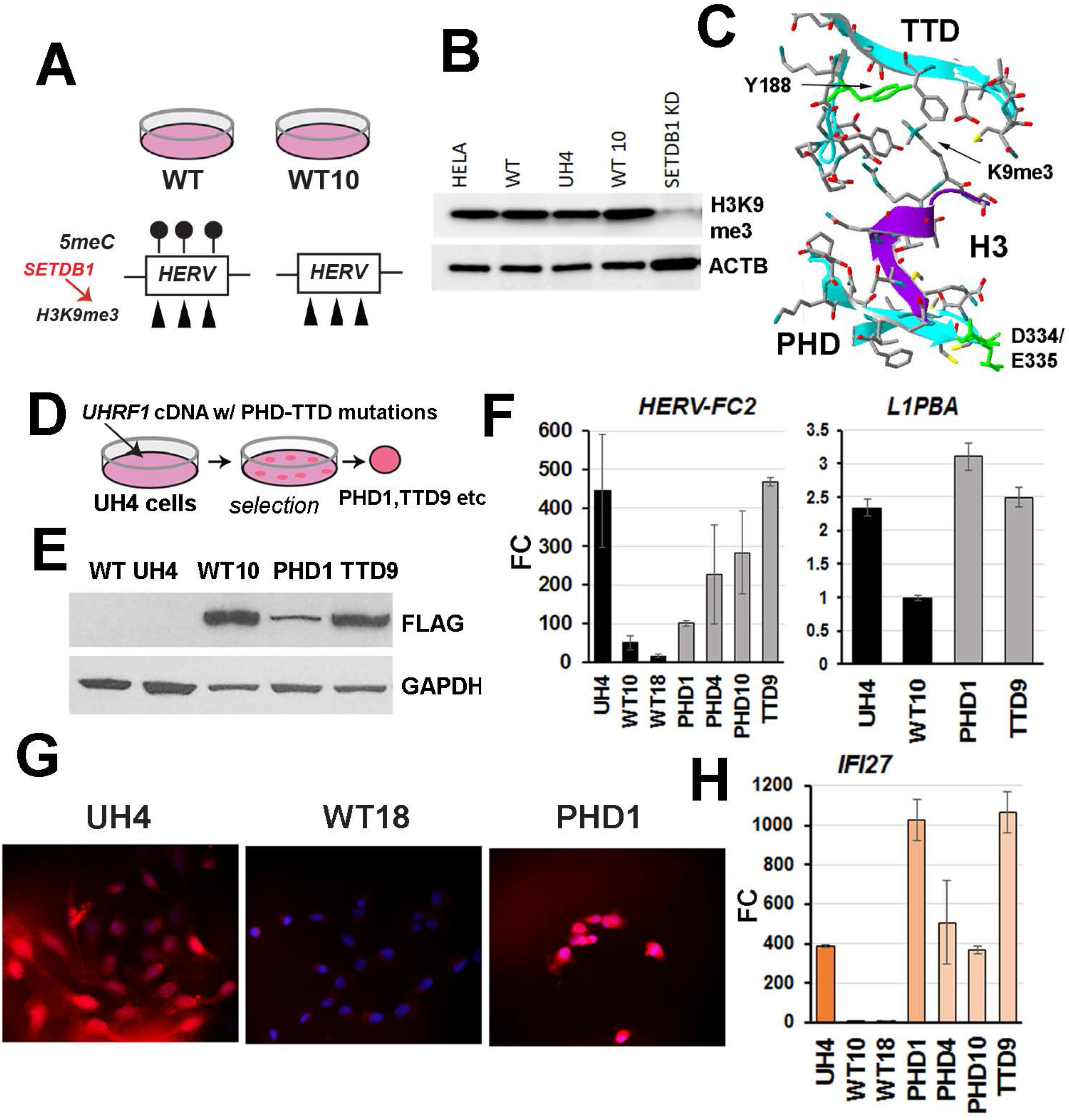
Knockdown cells cannot be rescued with UHRF1 proteins containing mutations known to affect H3K9me3 binding. (A) Assessing other mechanisms apart from DNA methylation which might contribute to HERV silencing in WT and rescued (WT10) cells. SETDB1 adds H3K9me3 to repress HERVs. (B) Knockdown of SETDB1 shows decrease in H3K9me3 levels than in normal cells (WT). (C) Model of the paired PHD-TTD domain of UHRF1 interacting with H3K9me3, showing the location of point mutations in the PHD (D334, E335) and TTD (Y188) previously shown to affect binding. (D) UH4 cells were transfected with cDNA as before, but containing the point mutations, and colonies expanded. (E) Western blot testing for FLAG-tagged proteins. (F) RT-qPCR of individual ERVs shows that re-establishment of repression does not occur to the same extent in cells rescued with UHRF1 proteins containing point mutations in PHD-TTD (PHD1, PH4, TTD9) as in those rescued with intact protein (WT10, WT18); error bars represent SEM. (G) Rescuing UH4 cells (top) with UHRF1 protein caused a shut-down of dsRNA production (middle-WT18) as detected by J2 antibody (red), but not if the protein contained point mutations affecting H3K9me3 binding (bottom-PHD1); nuclei were counterstained with DAPI (blue). (H) ISG response to dsRNA is still seen in UH4 cells rescued with point-mutated UHRF1 (PHD1, PHD4, PHD10, TTD9), but not in cells rescued with intact protein (WT10, WT18).

### Mutations in the PHD domain of mouse UHRF1 cause hypomethylation and transcriptional derepression of ERV in developing embryos

While our results so far implied that UHRF1 can potentially bind the H3K9me3 mark on ERV leading to repression of transcription from these elements, we did not see marked *de novo* methylation of the retroviruses in either the stable (Fig.5,6) or transient (Fig.4) experiments in hTERT1604. Since we have previously shown de novo methylation activity in these cells is sufficient to restore methylation to WT at some genes (O’Neill *et al*., 2018), we considered that these adult cells may instead lack other factors required for ERV DNA methylation which are only found earlier in development. To examine the dependence of *de novo* methylation and ERV repression on an intact H3K9me3 binding domain in UHRF1, we generated mouse embryos containing mutations in the PHD domain matching those used in human (Suppl.Fig.2A). To do so, we crossed B6D2F1 mice to generate zygotes, which we then injected with a single-guide RNA targeting the region around the DE amino acids in the PHD domain, together with an oligo containing the desired replacement nucleotides as well as an mRNA for the Cas9 enzyme. The first round of injections (306) and embryo transfers (11) resulted in no pups, suggesting that mutations were leading to embryonic lethality (Suppl.Fig.2A).

Consistent with this, homozygous mutant embryos (Uhrf1^DEAA/DEAA^) harvested at embryonic days 8.75 (E8.75) showed developmental delay and hypomethylation of IAP compared to WT (Suppl.Fig.2B). Both the retardation and the hypomethylation were more severe by E9.5 compared to WT (WT) or heterozygous (Uhrf1^WT/DEAA^) embryos (Suppl.Fig.2B, lower panels). A further round of injection gave one heterozygous founder animal (#13), which was back-crossed for one generation before intercrossing the heterozygous offspring to generate litters containing all three genotypes (Suppl.Fig.2A). Examination of DNA methylation at IAP 5’UTR showed highly significant decreases in homozygous embryos (HOM) compared to WT or heterozygous (HET) littermates (Suppl.Fig.2B). This decreased methylation was seen across the whole promoter region assayed. Highly significant decreases in DNA methylation compared to WT was also seen at other ERV regions assayed (data not shown). Whole mount *in situ* hybridisation for *IAP* showed strong homogenous signalling at e9.5 for homozygous mutant embryos (Fig.7A), with corresponding loss of DNA methylation at both the IAP LTR and 5’UTR regions (Fig.7B). We also examined transcription of ERV in the HOM embryos. These showed significant derepression of IAP and musD ERV as assayed by RT-qPCR (Fig. 7C), although increases were very variable across individual embryos. Analysis of both ERV and ISG transcription results from RT-qPCR confirmed that retroviral elements belonging to several classes, as well as interferon alpha and a number of ISG, were all generally more active in HOM mutants than in HET or WT (Fig.7C,D). In contrast, *Uhrf1* mRNA levels were more even across all embryos assayed (Fig.7C).

**Figure 7.**
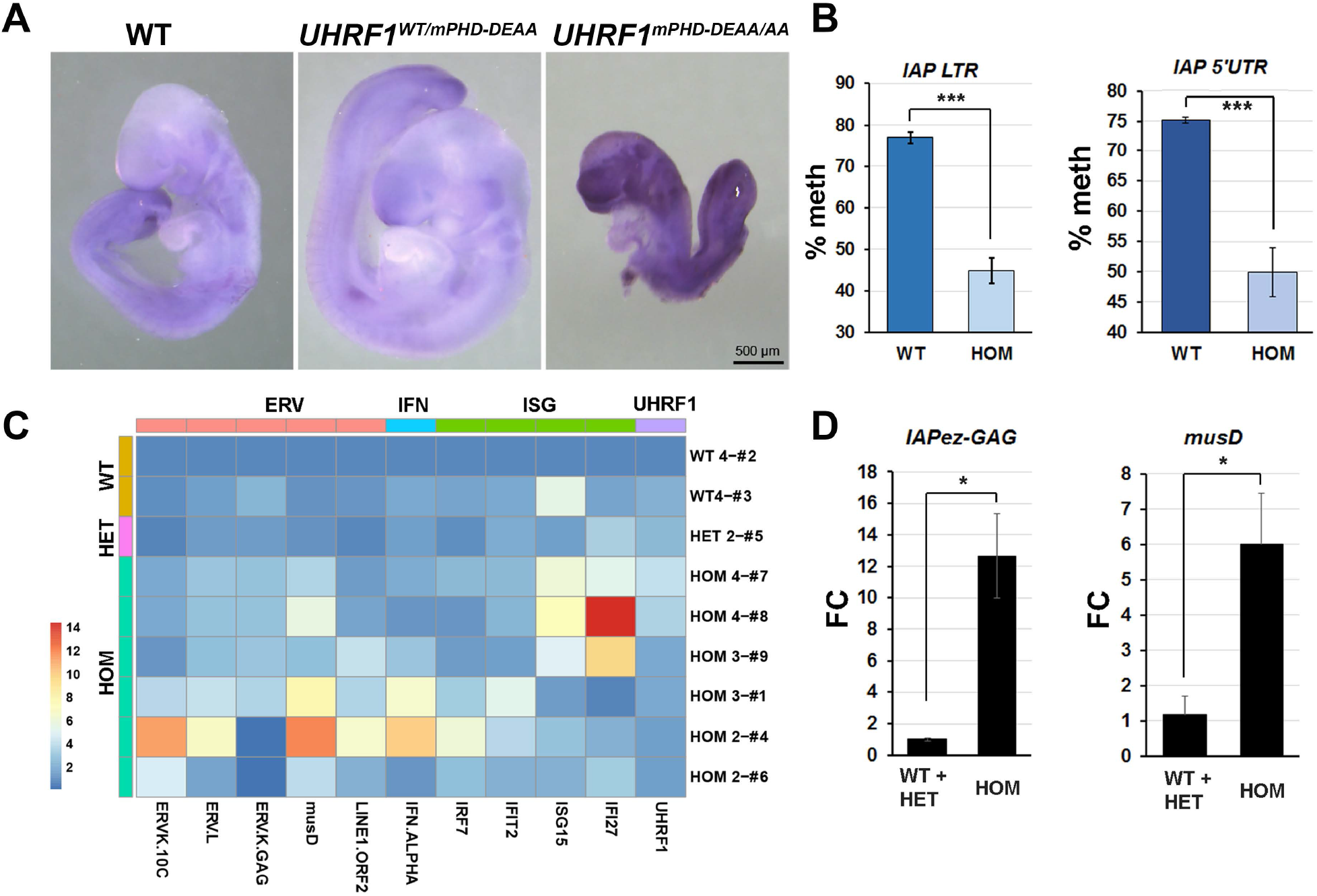
A conserved role for the PHD domain of UHRF1 in ERV methylation and repression in mouse. (A) Whole-mount in situ hybridisation on wild-type (WT), heterozygous (Uhrf1^WT/mPHD-DEAA^) and homozygous mouse embryos (Uhrf1^mPHD-DEAA/DEAA^) at E9.5 probed for IAP. Homozygous mutants showed developmental delay and IAP transcript was detected was throughout the embryo. Heterozygotes were indistinguishable morphologically from WT. (B) Overall methylation (% meth) at the IAP LTR is lower, as assessed by pyroassay, in homozygous mutant embryos (HOM); error bars represent SEM, results representative of at least 3 different embryos. (C) Heat map showing differences in expression of the indicated ERV, IFN and ISG (top) across embryos of the genotypes shown at left. Individual embryos and genes are indicated at right and bottom, respectively. *Uhrf1* mRNA levels are indicated at right as a control. Variation among F2 individuals may in part be due to segregation of different alleles. (D) RT-qPCR of individual ERVs showed significant derepression in homozygous mutant (HOM) embryos compared to WT.

**Figure 8.**
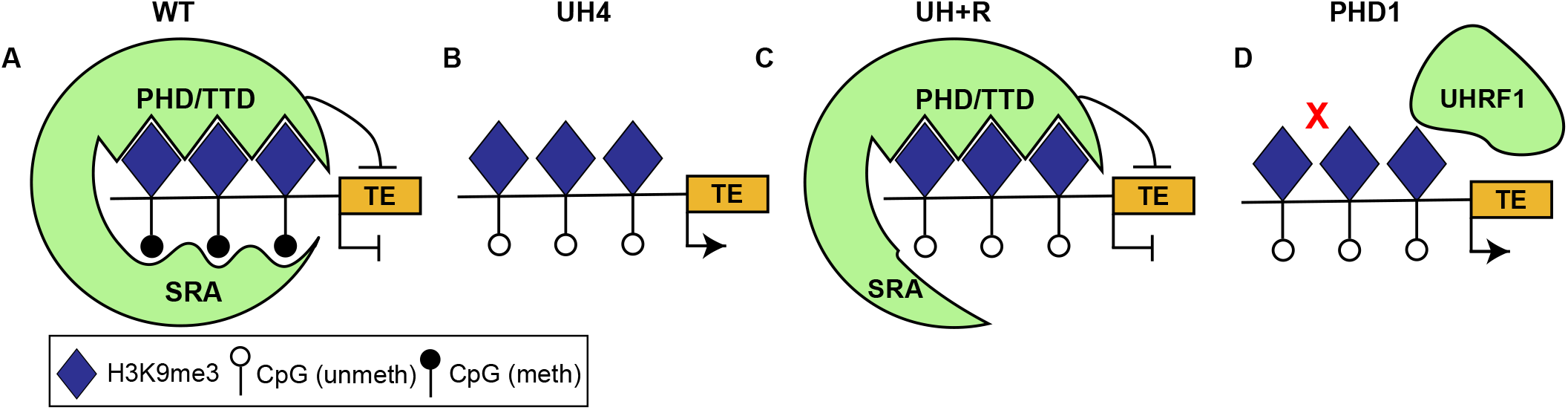
Graphical model depicting the role of UHRF1 in ERV suppression in adult cells. (A) In normal adult cells, UHRF1 binds to H3K9me3 through its PHD/TTD domain and to hemimethylated DNA through its SRA domain, recruiting DNA to methylate both DNA strands and suppressing ERV transcription. (B) In the absence of UHRF1, such as in the UH4 cell context, passive hypomethylation results in expression of ERV. (C) Rescue of UHRF1 to depleted cells enables ERV suppression, even in absence of DNA methylation at ERVs. (D) Functional mutation at the PHD domain of UHRF1 results in failure to suppress ERV expression.

## Discussion

We showed here that depletion of UHRF1 protein in differentiated human cells, either transiently or using stable models, causes loss of DNA methylation and an up-regulation of ERVs and a viral mimicry response (Chiappinelli *et al*., 2015; Roulois *et al*., 2015) from the cells, where genes involved in the innate immune response to viral infection, as well as CTA and other response genes, are markedly up-regulated. We show that this is linked to the presence of dsRNA in the cytoplasm, likely originating from de-repressed ERV elements, since in rescued cells where the ERV have been silenced, the dsRNA disappeared. Notably, this rescue effect can occur without reintroducing DNA methylation, suggesting a separate mechanism for ERV repression independent of the former but still dependent on UHRF1. Mutation in the PHD/TTD domain strongly implicated H3K9me3 as the signal which allowed UHRF1 to bind to ERV and repress them, and knockdown work suggested that the transcriptional co-repressor KAP1 is the mechanism by which UHRF1 affects this silencing. Consistent with this, mutating the histone binding domain of UHRF1 in mouse prevented the protein from recruiting DNA methylation to mouse ERVs post-implantation, resulting in widespread derepression, an innate immune response and embryonic lethality.

While the UHRF1 protein has been implicated in various different processes in the cell, including cell cycle regulation (Muto *et al*., 2002), DNA damage repair (Muto *et al*., 2002) and regulation of DNMT1 via ubiquitination (Qin *et al*., 2015) among others, a striking feature of the depletion studies carried out here is that the transcriptional response to the loss of this protein is dominated by a single process, viral mimicry. Thus, all ten of the top enriched GO terms in hTERT1604 fibroblasts stably depleted of UHRF1 were part of this response, including up-regulation of innate immune signalling, CTA derepression and antigen presentation. While there was less marked enrichment overall in HCT116 cells analysed from a prior study (Cai *et al*., 2017), 4/7 of the top terms were relevant to viral mimicry. We also found clear evidence of activation of ERV and of innate immune genes in melanoma cells depleted of UHRF1.

While overall the response to the loss of UHRF1 is very similar to that seen with small molecule inhibitors of DNMT1 (Chiappinelli *et al*., 2015; Roulois *et al*., 2015) or on the mutation of SETDB1 (Cuellar *et al*., 2017), the pathway uncovered here does differ in some details from those reports. While the major transcription factor implicated in response to DNMTi was IRF7, we found stronger up-regulation of IRF9 and STAT1 in our fibroblast cells. It may be that IRF9 is more important in our fibroblast lines than in the epithelial cell lines used previously for the array. While both this study and previous ones used JAK inhibitors to show effective blockade of the downstream signalling, the relative importance of the two IRF proteins in each cell type remains to be established. We also found some evidence for novel immune cell attractants not identified in the previous work which were upregulated in response to the loss of UHRF1, including the cell surface molecules CD3D and LY6E and the secreted cytokine CCL5. Some or all of these, in combination with interferons, may contribute to the extra-cellular signalling seen on the exposure of untreated cells to conditioned media from UHRF1 knockdown cells. While it is beyond the scope of the current work, it would be valuable to explore the relative roles of the various signals in attracting immune cells in an *ex vivo* or *in vivo* model, as well as to elucidate the details of the intracellular signalling pathway, particularly in comparison to DNMTi.

The initial knockout of *Uhrf1* in mouse also reported a widespread loss of DNA methylation and concurrent derepression of *LINE1* and *IAP* (Sharif *et al*., 2007), though in the absence of genome-wide methylation and transcription data, the main transcriptional response was unclear. A later paper by the same group also showed some up-regulation of ERV, but this was less than that seen in mice hypomorphic for *Dnmt1* (Sharif *et al*., 2016), where robust *IAP* upregulation had been originally reported (Walsh, Chaillet and Bestor, 1998), leading Sharif and colleagues to conclude that *Dnmt1* played a more important role in ERV repression than *Uhrf1*. They did not look at the transcriptional response for non-viral genes in response to either *Dnmt1* or *Uhrf1* knockout, and so did not uncover any ISG activation. In contrast, we found that in stable hypomorphic human fibroblast cells, the viral mimicry response was much stronger for *UHRF1* knockdowns than *DNMT1*. This is likely to be due to the greater sensitivity of cells to loss of DNMT1 protein, which triggers an effective DNA damage response (DDR) in a cell-autonomous fashion (Jackson-Grusby *et al*., 2001; Chen *et al*., 2007; Loughery *et al*., 2011; Liao *et al*., 2015), leading to the removal of cells with low levels of the protein and a strong selection for those which remethylate the DNA. In contrast, *UHRF1* mutations appear to be better tolerated as evidenced by recovery of knockdown clones at greater frequency and the more widespread and greater DNA demethylation seen in *UHRF1* cells (this study). Consistent with this, Meissner and colleagues reported a cell-autonomous death in normal human ES cells on deletion of *DNMT1* (Liao *et al*., 2015), and Cai et al have reported greater success in demethylating the genome by depleting *UHRF1* than using DNMTi (Cai *et al*., 2017). The greater DDR signalling effect in *DNMT1* hypomorphs (Loughery *et al*., 2011; Liao *et al*., 2015) may explain why only some features of viral mimicry have been reported in *DNMT1* hypomorphic colon cancer cells (Chiappinelli *et al*., 2015), and why only a partial signature (CTA genes activation) can be detected in our *DNMT1* hypomorphs (O’Neill *et al*., 2018).

In human cell lines, Cai et al recently showed partial activation of an antiviral response, including some ISGs in response to stable knockdown of *UHRF1* in HCT116 cells (Cai *et al*., 2017), but did not look at ERV methylation or transcription, or clearly identify the signalling pathways involved in response to ERV reactivation. Here, by reanalysing their array data, we could show that viral mimicry was the major transcriptional response in those cells, with CTA and type I interferon response genes being the major GO groups which were up-regulated.

In keeping with these observations, a recent analysis by Sadler and colleagues of transcriptional response in zebrafish to mutations in *Uhrf1* reported that the main effects seen were upregulation of innate immune and cell death signalling pathways (Chernyavskaya *et al*., 2017). Exploration of this system uncovered demethylation of ERV and a triggering of the innate immune response. In contrast to our results, they found both dsDNA and dsRNA signalling in the embryos, and favoured signalling through STING and cGAS rather than MDA5 and RIG1. In their case, an activation of DDR may have resulted in the generation of dsDNA as cells are killed and cleared from the embryo, which may explain the involvement of STING. In our case, 1) there was no evidence of up-regulation of any components of the dsDNA-sensing machinery; 2) the response we saw could be mimicked by exposing the cells to dsRNA instead; 3) dsRNA was directly detected in the RNA of *UHRF1* stable KD lines, but not WT; and 4) the dsRNA disappeared in U4 cells rescued with WT (but not mutated) UHRF1. However formal exclusion of a role for viral dsDNA will require further confirmatory work.

While some previous work on UHRF1 has highlighted a de-repression of ERV in response to loss or mutation of the protein (Sharif *et al*., 2007, 2016; Ramesh *et al*., 2016; Chernyavskaya *et al*., 2017), in all of these systems repression has been linked to DNA methylation of the ERV. Here we show for the first time that UHRF1 can suppress ERV expression in the absence of DNA methylation. This is borne out by 1) the reestablishment of transcriptional repression of ERV as UHRF1 levels recover in cells transiently depleted of the protein in the absence of any marked gain in DNA methylation; 2) the repression of ERV transcripts in stable knockdown cell lines rescued with WT UHRF1 protein (WT10), where DNA methylation levels remained low; 3)the disappearance of dsRNA signal in the WT10 cells; and 4) the switch-off of innate immune signalling in both transient and stable systems when UHRF1 levels are restored. SETDB1, which is a known repressor of ERV in mouse embryonic cells, was recently shown to be important for ERV repression in adult human leukaemia cell lines (Cuellar *et al*., 2017), where many of the same ERV were up-regulated as seen in our work. By using a small molecule inhibitor, we showed that SETDB1 was important for suppressing retroviral elements in our hTERT1604 cells too, and that cells depleted of UHRF1 were particularly sensitive to this inhibitor. SETDB1 trimethylates the H3K9 residue, a repressive mark commonly found on ERV, and these results, together with previous literature, suggested that H3K9me3 and DNA methylation both contribute to repression of ERV. We reasoned that retention of H3K9me3 in cells lacking DNA methylation might act as an instructive signal for UHRF1 allowing recognition of ERV when reintroduced. To test this, we used UHRF1 mutants lacking vital residues required to bind H3K9me3 (Rothbart *et al*., 2012, 2013) and found that in human cell lines, there was a failure to repress ERV, demonstrated by the presence of retroviral transcripts, dsRNA in the cytoplasm, and an innate immune response. Further, we showed that mice containing one of the same mutations failed to repress ERV *in vivo* and died mid-gestation. These results strongly suggest that UHRF1 binding to H3K9me3 on ERV through its TTD-PHD domain is required in vivo to repress ERV and that this is an important evolutionarily conserved role for the protein.

Our results from human cell lines and mouse imply that H3K9me3 alone is not sufficient to completely repress ERV, and that UHRF1 binding to this mark is required to ensure silencing and prevent activation of a viral response. In mouse, mutation of histone binding domain prevented silencing and embryos also showed a failure to gain DNA methylation at these elements post-implantation, suggesting that DNA methylation is an important component of long-term suppression, as we have previously shown in DNMT1 hypomorphs (Walsh, Chaillet and Bestor, 1998). However, in our adult human cell line experiments, it is clear that suppression is affected at least in part without the need for concomitant DNA methylation. The transcriptional corepressor KAP1 has been previously identified in mouse as an important suppressor of ERVs in ESC and preimplantation embryos (Matsui *et al*., 2010). Work from Trono and colleagues demonstrated that KAP1 was recruited to ERV by the KRAB-ZNF family of transcription factors (Wiznerowicz *et al*., 2007) which appear to coevolve with ERV and are important for their suppression (Ecco *et al*., 2016; Tie *et al*., 2018). Further, the recruitment of both SETDB1 and DNA methyltransferases appears to depend in part on KAP1 (Wiznerowicz *et al*., 2007; Ecco *et al*., 2016). Several sources have indicated that SETDB1-mediated suppression is important pre-implantation, with DNA methylation playing a greater role post-implantation in mouse (Wiznerowicz *et al*., 2007; Matsui *et al*., 2010; Ecco *et al*., 2016). Recently, a role for KAP1 in adult human cells has been established (Tie *et al*., 2018): we also found that depletion of SETDB1 caused derepression of ERV, and further could show that cells lacking UHRF1 and DNA methylation were particularly sensitive to loss of SETDB1. Our experiments suggest that UHRF1 may act as the link between H3K9me3 and DNA methyltransferase recruitment to ERV, particularly in the context of differentiating tissues, since while DNA methylation was present on ERV in our mouse model at e8.75, this remained static and did not increase to e9.5. While this needs further exploration, it is nevertheless clear that SETDB1 and H3K9me3 alone are not sufficient to prevent ERV derepression and activation of an innate immune response in either differentiated human or mouse cells, and that an intact UHRF1 protein is necessary in both systems. While the SRA domain of UHRF1 has been conclusively shown to bind hemi-methylated DNA (Bostick *et al*., 2007; Qin *et al*., 2015) and UHRF1 as a whole is known to be required for recruitment of DNMT1 to the replication fork (Bostick *et al*., 2007; Sharif *et al*., 2007), it is clear that without an intact TTD-PHD, SRA-mediated binding alone is not sufficient to abrogate ERV activation or immune signalling.

In conclusion, we have shown here that the major transcriptional response evoked by the loss of the UHRF1 protein was one of viral mimicry, including derepression of ERV and activation of a type I interferon response. An intact UHRF1 protein could re-establish repression on ERV even in the absence of DNA methylation, suggesting an independent mechanism for transcriptional suppression which relied on an intact H3K9me3 binding pocket in the protein. Mutations in this pocket prevented the suppression of ERV in human and mouse, and in the latter also affected DNA methylation patterns on ERV.

## Materials and Methods

### Cell culture

The wild-type (hTERT1604) lung fibroblast cell line (Ouellette *et al*., 2000) and derivatives were cultured in 4.5g/l glucose DMEM with 10% FBS and 2× NEAA (all Thermo-Fisher Scientific, Loughborough, UK). SK-MEL-28 cells (kind gift of Dr. Paul Thompson) were cultured in 4.5g/l glucose DMEM supplemented with 10% FBS. The hTERT1604 cell lines stably depleted of DNMT1 have been previously described (Loughery *et al*., 2011). Depletion of UHRF1 in hTERT1604 (U5/U10/UH4 lines) for this study used GIPz Lentiviral shRNAmir (Thermo-Fisher Scientific) according to the manufacturer’s instructions. Briefly, overlapping primers incorporating siRNA sequences to target UHRF1 were made and ligated into pGIPz.

The vector was linearized using *XhoI* and *MluI*, then 1□g transfected into WT cells using Lipofectamine 2000 (Thermo-Fisher Scientific) prior to selection in puromycin (Sigma-Aldrich, Dorset, UK) to isolate single colonies, which were then expanded; selection was removed 24 hours (24hrs) prior to any experimental analysis (Suppl. Table 1). Rescue cell lines (UH+R10/18, PHD1/4/10, TTD9) were generated by transfecting UH4 cells with pCMV plasmids containing full length UHRF1 cDNA which was either intact (WT) or contained functional mutations in either the PHD or TTD domains as previously described (Rothbart *et al*., 2013); individual colonies were selected in G418 (Sigma-Aldrich) and expanded as above. For transient knock-down experiments, 1×10^6^ cells/well were seeded in 6-well plates prior to reverse transfection basically as before (O’Neill *et al*., 2018) using 100nM ON-TARGETplus SMARTpool siRNA (Suppl. Table 2) or scrambled control (all ThermoFisher Scientific). Posttransfection, cells were cultured in complete medium to allow recovery, with extraction of RNA and DNA up to 28 days after addition of siRNA.

For drug treatment, Ruxolitinib (Absource, München, Germany) was dissolved in DMSO and added to culture media at a final concentration of 2□M; negative controls contained DMSO only (Suppl. Table 8). For analysis of dsRNA and dsDNA sensing pathways, cells were treated at a final concentration of 10μg/ml Poly(I:C) or sonicated salmon sperm DNA (Agilent, Stockport, UK) for 72hrs, with fresh media and drug every 24hrs; the nucleic acids were dissolved in sterile phosphate-buffered saline (PBS), heated at 50°C and cooled on ice to achieve re-annealing into double strands prior to treatment. For the media transfer test, UH4 cells were seeded and grown for 72hrs, then media transferred onto the wild type hTERT1604 cells, which were grown for another 72hrs.

### J2 staining

Cells were seeded onto glass slides pre-sterilized with 100% ethanol and UV light and allowed to attach overnight. Cells were fixed in 4% paraformaldehyde in PBS, 10mins before quenching with 0.1M glycine (Sigma-Aldrich), then permeabilized with 0.1% Triton X-100, 15mins and preblocked with 2% BSA (Sigma-Aldrich) for 1hr, room temperature (RT). Slides were incubated with J2 primary antibody (Scicons, Szirák, Hungary) at 1:200 in 2% BSA overnight at 4°C. The next day, slides were washed and incubated with anti-mouse IgG AlexaFluor 546 (Invitrogen, Paisley, Scotland) antibody at 1:1000, 1hr, RT before washing and adding DAPI mounting media (Santa Cruz Biotechnology, Heidelberg, Germany) (Suppl. Table 3). Fluorescent images were taken with a Nikon Eclipse E400 phase contrast microscope and processed using Adobe Photoshop (Maidenhead, UK).

### DNA analyses

Genomic DNA was extracted from cells growing in log phase, with each cell line done in triplicate, including one biological replicate. DNA preparation, bisulfite conversion and array hybridization were essentially as previously described (Mackin, O’Neill and Walsh, 2018; O’Neill *et al*., 2018). Briefly, DNA was isolated using the QIAmp DNA Blood Mini Kit (Qiagen, Crawley, UK), assessed for integrity and quality using a range of measures including agarose gel electrophoresis, UV absorbance and Quant-iT PicoGreen dsDNA assay (Thermo Fisher Scientific). Purified DNA was sent to Cambridge Biological Services where bisulfite conversion was performed using the EZ DNA Methylation kit (Zymo Research, California, USA) and samples were loaded onto the Infinium HumanMethylation450 BeadChip (Bibikova *et al*., 2011) and imaged using the Illumina iScan.

Genomic DNA was extracted from mouse tail as previously described (Rutledge *et al*. 2014). Genotyping was performed on lysate using primers listed in Suppl. Table 4.

For pyrosequencing, DNA (500ug) was bisulfite-converted in-house as above, then PCR-amplified using the PyroMark PCR kit using Qiagen’s pyrosequencing primer assays or those designed in-house (Suppl. Table 5) via the PyroMark Assay Design Software 2.0 (Qiagen). Reaction conditions were as follows: 95°C, 15mins; followed by 45 cycles of 94°C, 30secs; 56°C, 30secs and 72°C, 30secs; final elongation 72°C, 10mins, with products verified on agarose gels prior to pyrosequencing using the PyroMark Q24 (Qiagen).

### RNA analyses

RNA was extracted from cells growing in log phase using the RNEasy Mini Kit (Qiagen, Crawley, UK), including a DNAse step, according to manufacturer’s instructions. Complementary DNA (cDNA) was reversed transcribed in a reaction containing 250-500ng total RNA, 0.5uM dNTPs, 0.25ug random primers (Roche, UK), 1x reverse transcriptase buffer and 200U RevertAid reverse transcriptase in a total volume of 20ul. Reaction conditions were as follows: 25°C, 10 minutes (mins); 42°C, 60mins; 70°C, 10mins. cDNA was stored at −80°C until use. Each RT-PCR reaction contained 1ul cDNA from the above reaction, 1x buffer, 0.4mM dNTPs, 1uM primers (Suppl. Table 6), MgCl_2_ concentration specific to the primers and 0.01 U Taq polymerase. Reaction condition were as follows: 94°C, 3mins; followed by cycles of 94°C, 30 seconds (secs); gene-specific annealing temperature for 1min; 72°C, 1min; with final elongation at 72°C, 5mins. RT-qPCRs were performed using 1× LightCycler 480 SYBR Green I Master (Roche), 0.5 μM primers (Suppl. Table 7) and 1 μl cDNA. Reactions were run on the LightCycler 480 II (Roche), with an initial incubation step of 95°C, 10mins; followed by 50 cycles of 95°C, 10secs; 60°C, 10secs and 72°C,10 secs. Expression was normalised to *HPRT*, and relative expression was determined using the ΔΔC_T_ method.

Array work was carried out essentially as previously described (Mackin, O’Neill and Walsh, 2018; O’Neill *et al*., 2018): briefly, total RNA was extracted from each cell line growing in log phase in triplicate, including at least one biological replicate, and was assessed for integrity and quantity using a SpectroStar (BMG Labtech, Aylesbury, UK) and bioanalyser (Agilent Technologies, Cheadle, UK) prior to sending to Cambridge Analytical Services for linear amplification using the Illumina TotalPrep RNA Amplification Kit (Life Technologies/Thermofisher, Paisley, UK) followed by hybridization to the HumanHT-12 v4 Expression BeadChip.

### Protein analyses

Cells growing in log phase were harvested for protein extraction using the protein extraction buffer (50 mM Tris–HCl, 150 mM NaCl, 1% Triton-X, 10% glycerol, 5 mM EDTA; all Sigma-Aldrich) and 0.5 μl protease inhibitor mix (Sigma-Aldrich). For Western blotting, 30μg protein was denatured at 70°C in the presence of 5μl 4× LDS sample buffer and 2μl 10× reducing agent (Invitrogen) in a total volume of 20μl nuclease-free water (Qiagen). Proteins were separated by SDS-PAGE and electroblotted onto a nitrocellulose membrane (Thermo-Fisher Scientific), then blocked in 5% non-fat milk for either 1h at RT or overnight at 4°C. Membranes were incubated with the primary antibody overnight (Suppl. Table 3) overnight at 4 °C, followed by HRP-conjugated secondary antibody incubation at RT using ECL (Thermo-Fisher Scientific).

### Generation of the UHRF1 PHD D335/D336AA mutant mice

*B6D2F1* female mice (8-10 weeks old) were super-ovulated by intraperitoneal injection of pregnant mare’s serum gonadotropin (PMSG, 5 IU). Mice were injected with human chorionic gonadotropin (hCG, 5 IU) 48h later and mated with B6D2F1 males. Zygotes were collected from the oviducts of female mice at embryonic day (E) 0.5. The cumulus cells were removed by incubation in 1% hyaluronidase/M2 medium before washing with fresh M2 medium and recovery for 2-4h at 37 °C in a 5% CO_2_ incubator. SpCas9 mRNA (100 ng/μl), *Uhrf1*-sgRNA (10ng/μl), and 112-bp single-stranded oligodeoxynucleotide (ssODN, 50ng/μl), which was flanked by homologous arms corresponding to exon 7 of *Uhrf1 (Suppl. Table 9*), were mixed immediately before microinjection using a FemtoJet microinjector, set to Pc = 10-15 hPa, and Pi = 40-50 hPa. Successfully microinjected zygotes were incubated in KSOM at 37 °C in a 5% CO_2_ incubator for 24h until they reached the 2-cell stage and transferred into oviducts of 0.5 days post-coitum (dpc) pseudopregnant ICR females. To investigate CRISPR/Cas9-mediated mutation in the *Uhrf1* gene, genomic DNA was prepared from 3wk-old mouse tails. The genomic regions flanking the gRNA target were amplified by PCR using specific primers (Suppl. Table 4). The PCR amplicons were ligated into the pClone007 vector (Tsingke, TSV-007s) and sequenced.

### Whole mount *in situ* hybridisation

*In situ* hybridisation on whole embryos at e.95 was performed as previously described (Walsh, Chaillet & Bestor, Nat Gen 1998).

### Bioinformatics and statistical analysis

Output files in IDAT format were processed and bioinformatic analysis was carried using the RnBeads (Assenov *et al*., 2014) methylation analysis package (v1.0.0) as previously described (Mackin, O’Neill and Walsh, 2018). In order to map CpG sites showing highly reproducible changes (FDR <0.05) against the locations of RefSeq genes on the UCSC genome browser (Karolchik *et al*., 2003) for each cell line, we employed a bespoke workflow in the GALAXY platform (Giardine *et al*., 2005). Absolute *β* levels were used to measure median methylation across genes of interest using custom workflows in GALAXY, with further statistical analyses in Statistical Package for the Social Sciences software (SPSS) version 22.0 (SPSS UK Ltd).

Agilent arrays GSE93142 and GSE93135 from SuperSeries GSE93136 from (Cai *et al*., 2017) were processed using the R package GEOquery (2.46.15), annotation package hgug4112a.db (3.8) and annotation table for Agilent-014850 Whole Human Genome Microarray 4×44K G4112F (Probe Name version) from GEO to obtain log2 normalized fold changes (FC) per probe. Gene Ontology analysis through DAVID (da, Sherman and Lempicki, 2009) was then computed using the top 500 genes with greater than 1.5 FC.

Statistical analysis for pyrosequencing and RT-qPCR data were analysed by Student’s paired t-test using Microsoft Excel (Microsoft Office Professional Plus 2016). Experiments were carried out in duplicate and included at least one biological replicate. Error bars on all graphs represent standard error of the mean (SEM) or in the case of HT12 array data, 95% confidence interval (CI), unless otherwise stated. Asterisks are used to represent probability scores as follows: \**p* < 0.05; \*\**p* < 0.01; \*\*\**p* < 0.005 or n.s. not significant. Statistical analysis for the Illumina 450k methylation array was performed by the RnBeads package as previously described (Mackin, O’Neill and Walsh, 2018).

## Acknowledgments

We thank Paul Thompson for kind donation of SK-MEL-28 cells. We are obliged to Bert Vogelstein for the HCT116 WT and DKO cells. We thank other laboratory members for technical help, advice and critical comments.

---

**Supplementary Figure 1.**
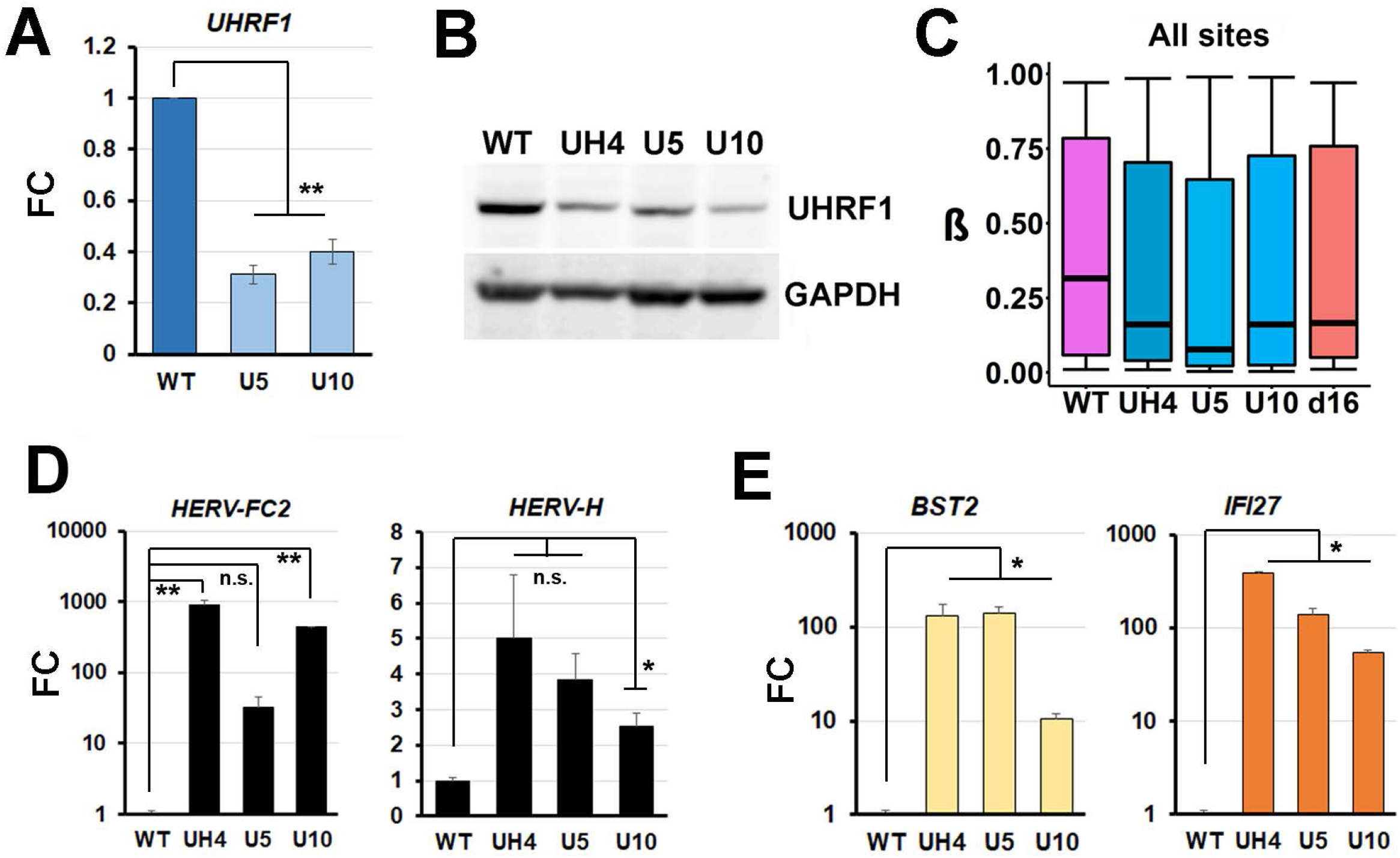
(A) RT-qPCR confirming *UHRF1* mRNA depletion in stable knockdown clones U5 and U10. (B) Western blot for UHRF1, GAPDH used a loading control. (C) Overall median methylation from 450k array for *UHRF1*-depleted cell lines compared to the most demethylated *DNMT1-*depleted line (d16). (D) RT-qPCR confirmation of *ERV* upregulation in all *UHRF1* clones. (E) Upregulation of ISG in all *UHRF1* stable knockdown cell lines, confirmed by RT-qPCR.

**Supplementary Figure 2.**
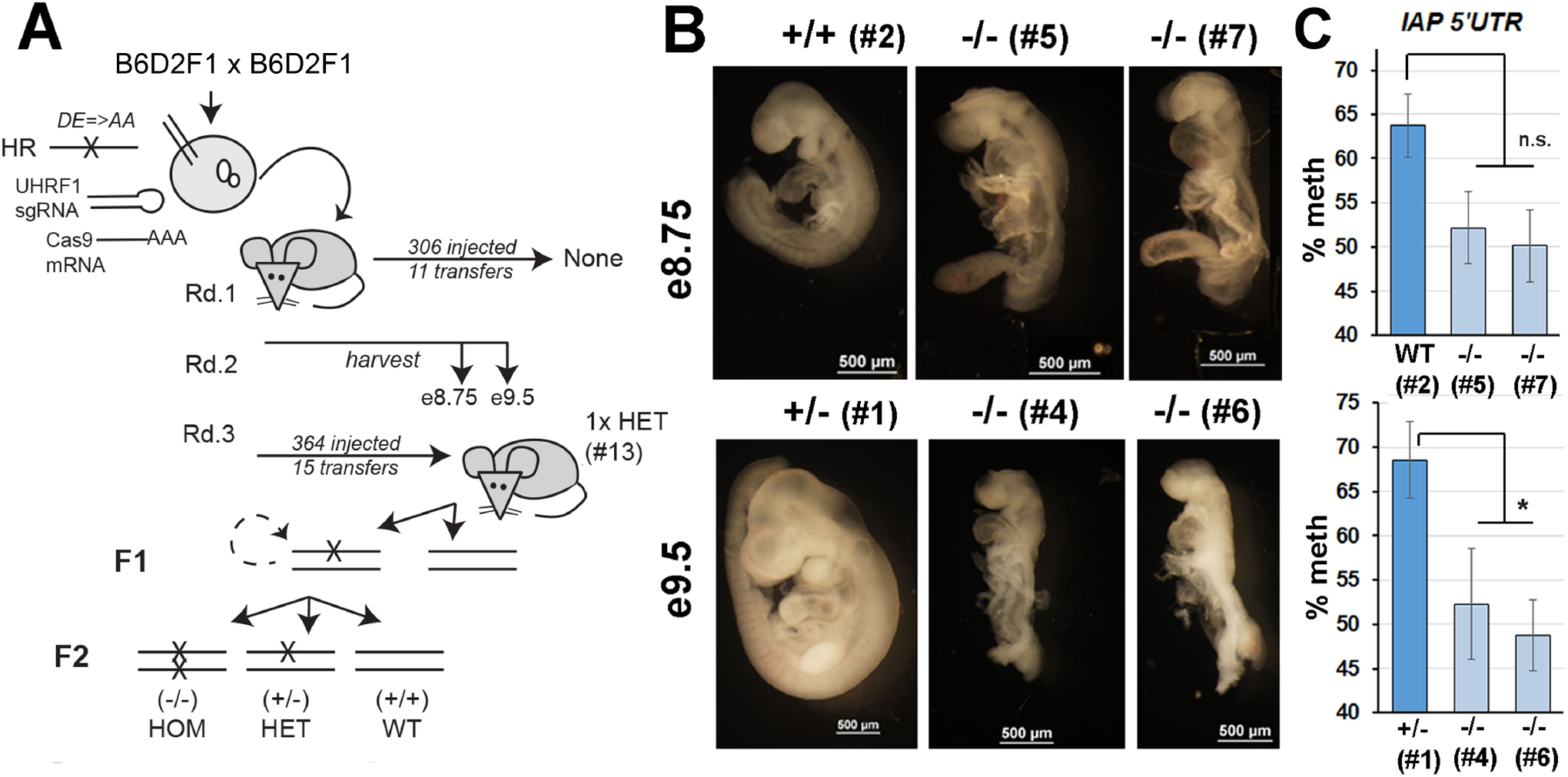
(A) Strategy used to generate mutant mice containing point mutations in the PHD domain. Inter-strain hybrid zygotes were injected with (i)a single-guide RNA for the targeted region in *Uhrf1*, (ii)a donor molecule for Homologous Recombination (HR) containing the mutations to match the D334A/E335A in human and (iii)an mRNA for Cas9. Round 1(Rd.1) of injections gave no mice, so pregnant females from Rd.2 were sacrificed at e8.75 and e9.5 to examine embryos (see B). Further injections (Rd.3) resulted in a single heterozygous (HET) pup (#13), who was back-crossed once, then offspring inter-crossed (F1) to generate homozygous (HOM) as well as heterozygous embryos. (B) Embryos from Rd.2 injections: at e8.75 homozygous mutant (-/-) mutants showed developmental delay (left) and differences in methylation at the 5’UTR of IAP elements by pyroassay in (C.) compared to WT (+/+). By e9.5 delay was more marked (left) and methylation differences significant (right). Heterozygotes were indistinguishable morphologically from WT at both e8.75 and e9.5 (e.g #1).

