## Supplementary Material for "UHRF1 suppresses viral mimicry through both DNA methylation-dependent and -independent mechanisms"

**Supplementary table 1. shRNA**

| Target | Supplier | Cat. number | Mature antisense sequence | Cell lines |
|---|---|---|---|---|
| ***UHRF1*** | Dharmacon | V3LHS_ 353720 | TGACATTGCGCACCACCCT | U prefix (U5, U10) |
| ***UHRF1*** | Dharmacon | V3LHS_413692 | AACGTTATATCTTTCTTGG | UH prefix (UH4, UH5) |

**Supplementary table 2. siRNA – ON-TARGETplus Human - SMARTpool**

| Target | Supplier | Cat. number |
|---|---|---|
| ***KAP1*** | Horizon Discovery | L-005046-00-0005 |
| ***SETDB1*** | Horizon Discovery | L-020070-00-0005 |
| ***UHRF1*** | Horizon Discovery | L-006977-00-0005 |
| ***MAVS*** | Horizon Discovery | L-024237-00-0005 |

**Supplementary table 3. Antibodies**

| Target | Supplier | Cat. number | Raised in | Clonality | Dilution | Size (kDa) | Application |
|---|---|---|---|---|---|---|---|
| **Primary antibodies** | | | | | | | |
| ***UHRF1*** | SC | 373750 | Mouse | Mono | 1:100 | 90 | WB |
| ***GAPDH*** | CST | 14C10 | Rabbit | Mono | 1:10000 | 36 | WB |
| ***FLAG*** | SA | F1804 | Mouse | Mono | 1:1000 | 120 | WB |
| **dsRNA** | SCICONS | 10010200 | Mouse | Mono | 1:200 | X | IF |
| **Secondary antibodies** | | | | | | | |
| **AR-IgG** | SC | sc-2004 | Goat | Poly | 1:10000 | X | WB |
| **AM-IgG** | SA | A9044 | Rabbit | Poly | 1:5000 | X | WB |
| **AM-IgG** | Invitrogen | A10036 | Donkey | Mono | 1:1000 | X | IF |

SC (Santa Cruz Biotechnology); CST (Cell Signalling Technologies); SA (Sigma-Aldrich); Mono (monoclonal); Poly (polyclonal); kDa (Kilodaltons).

**Supplementary Table 4: Genomic PCR primers**

| Gene | Primer | Oligo sequence (5’-3’) |
|---|---|---|
| ***Np95*** | FW | TGACTCTCAGCTCAACAACTGTCG |
| | RV | CTCTCACTGAGCCTACAGCCAAG |

**Supplementary Table 5. Pyrosequencing primers**

| Human pyrosequencing primers | | | |
|---|---|---|---|
| **Gene** | **Primer** | **Modification** | **Oligo sequence (5'-3')** |
| ***HERV-FC2*** | FW | | TTTGGTTTTTTTTGTTAGGATTAGTGA |
| | RV | biotin | AAATCTCCTCCCCAATTTTAACACCA |
| | SEQ | | TTTTTGTTAGGATTAGTGAATT |
| ***HERV-H*** | FW | | AGGGTTTGTGTGAGTAATAAAGTT |
| | RV | biotin | ACTCCTACCCCCCAAAAAACAAACT |
| | SEQ | | AGTTTTTAATTATTTGGGTGT |
| ***LINE-1*** | FW | | GGGAGGAGTTAAGATGGT |
| | RV | biotin | ATAAACCCCATACCTCAA |
| | SEQ | | GGGAGGAGTTGGATGGT |
| **Mouse pyrosequencing primers** | | | |
| **Gene** | **Primer** | **Modification** | **Oligo sequence (5'-3')** |
| ***Iap 5-UTR*** | FW | | GGGTTGTAGTTAATTAGGGAGTGATA |
| | RV | biotin | ACAATTAAATCCTTCTTAACAATCTACTT |
| | SEQ | | ATTTTGGTTTGTTGTGT |
| ***Iap LTR*** | FW | biotin | GGTTTTGGAATGAGGGATTTT |
| | RV | | CTCTACTCCATATACTCTACCTTC |
| | SEQ | | ATACTCTACCTTCCCC |
| ***Line-1*** | FW | | GTAGAAGTATAGAGGGGTTGAGGTA |
| | RV | biotin | ACAATTCCCAAATAATACAAACTCT |
| | SEQ | | AGTATTTTGTGTGGGT |

**Supplementary Table 6. RT-PCR primers**

| Gene | Primer | Oligo sequence (5'-3') |
|---|---|---|
| ***ACT-B*** | FW | GGACTTCGAGCAAGAGATGG |
| | RV | AGCACTGTGTTGGCGTACAG |
| ***UHRF1*** | FW | TGAGGACATGTGGGATGAGA |
| | RV | GTCCCTGGAGTTCATCTGGA |

**Supplementary Table 7. RT-qPCR primers**

| **Human RT-qPCR primers** | | |
|---|---|---|
| **Gene** | **Primer** | **Oligo sequence (5'-3')** |
| ***IFI27*** | FW | TCTGTCCACCCTCTGCTTCT |
| | RV | GGCATGGTTCTCTTCTCTGC |
| ***OAS2*** | FW | AGGAAGGTAGCGCATCTTGA |
| | RV | CACCTCTGTCCGACTTGGTT |
| ***STAT1*** | FW | CCCACTCTGATCAACTTTTGC |
| | RV | GGCCTGTTGAAGATGCTTGT |
| ***ISG15*** | FW | ACCTACGAGGTACGGCTGAC |
| | RV | GGTGGAGGCCCTTAGCTC |
| ***HERV-FC2*** | FW | TTTCCCACCGCTGGTAATAG |
| | RV | AGGCTAAGGATTCGGCTGAG |
| ***HERV-H*** | FW | TTCACTCCATCCTTGGCTAT |
| | RV | CGTCGAGTATCTACGAGCAAT |
| ***HERV-W1*** | FW | AAGAATCCCTAAGCCTAGCTGG |
| | RV | GCCTAATTAGCATTTTAGTGAGCTC |
| ***L1PBA*** | FW | CTTTGCAGACACTCCCCAGT |
| | RV | GGTCTAGCCACCCAGCAG |
| ***L1P1*** | FW | TGCCCTAAAAGAGCTCCTGA |
| | RV | TGTTTTTGCAGTGGCTGGTA |
| ***UHRF1*** | FW | TGAGGACATGTGGGATGAGA |
| | RV | GTCCCTGGAGTTCATCTGGA |
| ***HPRT*** | FW | AGCCCTGGCGTCGTGATTAGT |
| | RV | CCCGTTGAGCACACAGAGGCCTA |
| **Mouse RT-qPCR primers** | | |
| **Gene** | **Primer** | **Oligo sequence (5'-3')** |
| ***Gapdh*** | FW | CAACTACATGGTCTACATGTTC |
| | RV | CTCGCTCCTGGAAGATG |
| ***IAPez-gag*** | FW | CACGCTCCGGTAGAATACTTACAAAT |
| | RV | CCTGTCTAACTGCACCAAGGTAAAAT |
| ***MusD*** | FW | GAATATGTCTAATACGCTAGCCTTTCC |
| | RV | GTAATGTCTGCCCCTAGTATCTTGTT |
| ***Ifn-alpha*** | FW | CTGCTGGCTGTGAGGACATA |
| | RV | AGGAAGAGAGGGCTCTCCAG |

| | | |
|---|---|---|
| ***ERV-K10C gag*** | FW | ATGTGAGCTAGCTGTTAAAGAAGGAC |
| | RV | CTCTCTGTTTCTGACATACTTTCCTGT |
| ***ERV-K10C*** | FW | CCAAATAGCCCTACCATATGTCAG |
| | RV | GTATACTTTCTTCTTCAGGTCCAC |
| ***ERV-L gag*** | FW | TTCTTCTAGACCTGTAACCAGACTCA |
| | RV | TCCTTAGTAGTGTAGCGAATTTCCTC |
| ***Line1-ORF2*** | FW | GACATAGACTAACAAACTGGCTACACAAAC |
| | RV | GGTAGTGTCTATCTTTTTCTCTGAGATGAG |
| ***Ifi27*** | FW | TAACTGGTCCTCATGGCGTT |
| | RV | CCCCTTCGAACCAGCTAGAA |
| ***Uhrf1*** | FW | AAAACGCCCTGAGTTTTCGC |
| | RV | CCGATGTACTCTCTCACGGC |
| ***Isg15*** | FW | AGCAATGGCCTGGGACCTAA |
| | RV | AGACCCAGACTGGAAAGGGT |
| ***Ifit2*** | FW | CTGGGGAAACTATGCTTGGGT |
| | RV | ACTCTCTCGTTTTGGTTCTTGG |
| ***Irf7*** | FW | CCCATCTTCGACTTCAGCAC |
| | RV | TGTAGTGTGGTGACCCTTGC |

**Supplementary table 8. Pharmacological Inhibitors**

| Target | Supplier | Cat. number |
|---|---|---|
| **Ruxolitinib (INCB018424)** | Selleckchem | S1378 |

**Supplementary table 9. Mouse CRISPR guide/oligo sequences**

| Target | Oligo sequence (5’-3’) FW | Oligo sequence (5’-3’) RV |
|---|---|---|
| **sgRNA sequence** | | |
| ***UHRF1 exon 3*** | GTTGTGTGATGAGTGTGACA | AAAAAAGCACCGACTCGGTG |
| **PHD mutant HDR oligo sequence** | | |
| ***UHRF1 exon 7*** | GTGCCTGCCATGTGTGTGGTGGGCGCGAGGCTCCTGAGAAACAGCTGTTGT GTGCTGCGTGCGATATGGCCTTCCACCTGTACTGCCTGAAGCCACCGCTCA CCTCTGTCCC | |
